# Brain Network Reconfiguration for Narrative and Argumentative Thought

**DOI:** 10.1101/2020.07.20.211466

**Authors:** Yangwen Xu, Lorenzo Vignali, Olivier Collignon, Davide Crepaldi, Roberto Bottini

## Abstract

Our brain constructs reality through narrative and argumentative thought. Some hypotheses argue that these two modes of cognitive functioning are irreducible, reflecting distinct mental operations underlain by separate neural bases; Others ascribe both to a unitary neural system dedicated to long-timescale information. We addressed this question by employing inter-subject measures to investigate the stimulus-induced neural responses when participants were listening to narrative and argumentative texts during fMRI. We found that following both kinds of texts enhanced functional couplings within the frontoparietal control system. However, while a narrative specifically implicated the default mode system, an argument specifically induced synchronization between the intraparietal sulcus in the frontoparietal control system and multiple perisylvian areas in the language system. Our findings reconcile the two hypotheses by revealing commonalities and differences between the narrative and the argumentative brain networks, showing how diverse mental activities arise from the segregation and integration of the existing brain systems.

## Introduction

“To say that all human thinking is essentially of two kinds – reasoning on the one hand, and narrative, descriptive, contemplative thinking on the other – is to say only what every readers’ experience will corroborate.”

*William James*

Humans are thinking animals. Flows of concepts and ideas pass through our minds from time to time. These concepts and ideas are seldom in isolation; they are often sequentially connected, composed into a mental discourse, which has been called the “train of thought” (Hobbes, 1651). Psychologists argued for decades that these complex thoughts are essentially of two natural kinds, each gluing its elements in a different manner (Bruner, 1986; James, 1983): The narrative thought comprises a series of events, which unfold through temporal causality and implied purpose (Beach and Bissel, 2016). The argumentative thought consists of a chain of propositions, forming the interlinked premiss-illative-conclusion structure, according to which a final conclusion is reached through progressive inferences (Hitchcock, 2007).

Despite the fact that both modes of thought are pervasive in our mental life, most neuroimaging studies merely focused on the neural basis of narrative thought (Kemmerer, 2014; Mar, 2004). In these studies, a narrative text is divided into its constituent sentences, and the order of these sentences is randomized to form a sentence-scrambled version of the text. The conditions presenting the intact texts are contrasted to conditions presenting the sentence-scrambled texts. As participants can only generate a coherent narrative discourse in the intact-text condition, this contrast outstands the neural basis of narrative thought from the one of linguistic processing regarding word meaning and syntax. A meta-analysis of 12 such neuroactivation studies indicates that narratives consistently induced greater activation than the sentence-scrambled text in the anterior temporal lobe, temporoparietal junction, precuneus, and medial prefrontal cortex (Ferstl et al., 2008); a set of regions that coincides with the default mode network (DMN) (Buckner et al., 2008). Instead of investigating the overall level of activation, recent studies demonstrate that the DMN activity can also capture the dynamic progress in a narrative (Lerner et al., 2011; Simony et al., 2016). As it is hard to obtain an explicit event-related response model that can describe a narrative discourse, these studies used one individual’s neural response to model another’s by measuring the shared neural responses across participants when they were listening to the same narrative (Nastase et al., 2019). For instance, one study using the inter-subject correlation (ISC) method find that listening to the same narrative synchronizes the blood-oxygen-level-dependent (BOLD) fluctuations in the same regions of the DMN across subjects; listening to the same sentence-scrambled text does not (Lerner et al., 2011). Another study further illustrates such higher synchronization in the DMN not only exist between the same regions across subjects (i.e., ISC) but also between different regions across subjects (i.e., the inter-subject functional connectivity, ISFC) (Simony et al., 2016). The later findings demonstrate that regions in the DMN underlie narrative thought by coordinating with each other as a network.

What are the neural bases of argumentative thought? There are two hypotheses (Jacoby and Fedorenko, 2018). The content-dependent hypothesis inherits the two modes of thought view, suggesting the narrative and the argumentative thought are irreducible to one another (Bruner, 1986; James, 1983); they reflect distinct mental operations, which should correspond to separate neural bases. Tracing a narrative plot relies on constructing and updating the representation of a state of affairs, i.e., “situation model” (Zwaan and Radvansky, 1998), to simulate the temporal causality and to infer the characters’ intentions. This set of cognitive functions is indeed attributed to the DMN, which plays a role in mental simulation and theory of mind (Buckner et al., 2008). Following an argument, instead, relies on identifying and evaluating the logical structure embedded in the use of natural language, i.e., “informal logic” (Blair, 2015). This set of cognitive functions might warrant cooperation between the language and the reasoning brain system.

On the contrary, the content-independent hypothesis suggests that the narrative and the argumentative thought are fundamentally the same; they share the same neural mechanism. One commonality of these two modes of thought is that the content at each time point relates to the context established at previous time points. Iteratively accumulating information over time and holding the information online over a long timescale seems equally crucial to framing a coherent narrative and a valid argument. According to the hierarchical process memory framework, all the cortical circuits accumulate information over time, but their processing timescale increases along the hierarchical topography, from milliseconds in primary sensory regions to minutes in high-order regions (Hasson et al., 2015). This framework suggests that the DMN, which is at the top of the topographical hierarchy (Margulies et al., 2016; Sepulcre et al., 2012), supports narrative thought by virtue of its wide temporal receptive window (TRW), integrating information over a long timescale, e.g., up to minutes (J. Chen et al., 2016). As a wide TRW is also crucial to the progress of an argumentative thought, the DMN might potentially serve as general machinery for long-timescale information integration, supporting both narrative and argumentative thought.

Testing these two hypotheses requires to fill the vacancy of studies on argumentative thought. Here, we investigated the neural correlates of both narrative and argumentative thought by contrasting the BOLD signal elicited by two narrative texts and two argumentative texts to the signal elicited by their corresponding sentence-scrambled version (Table 1). We also acquired the BOLD signal during the resting state as a baseline. Specifically, we employed ISC and the ISFC as measures to respectively investigate the stimulus-induced regional activity and interregional functional coupling during the narrative and the argumentative thought. The content-independent hypothesis will predict a higher ISC or ISFC in the DMN in both the narrative and the argumentative conditions compared to their corresponding sentence-scrambled conditions. The content-dependent hypothesis, instead, will predict a higher ISC or ISFC in the DMN only when the narrative condition and its sentence-scrambled condition are compared; alternative brain networks that relate to language and reasoning will engage in the discourse-level comprehension of argumentative texts.

**Table 1.**
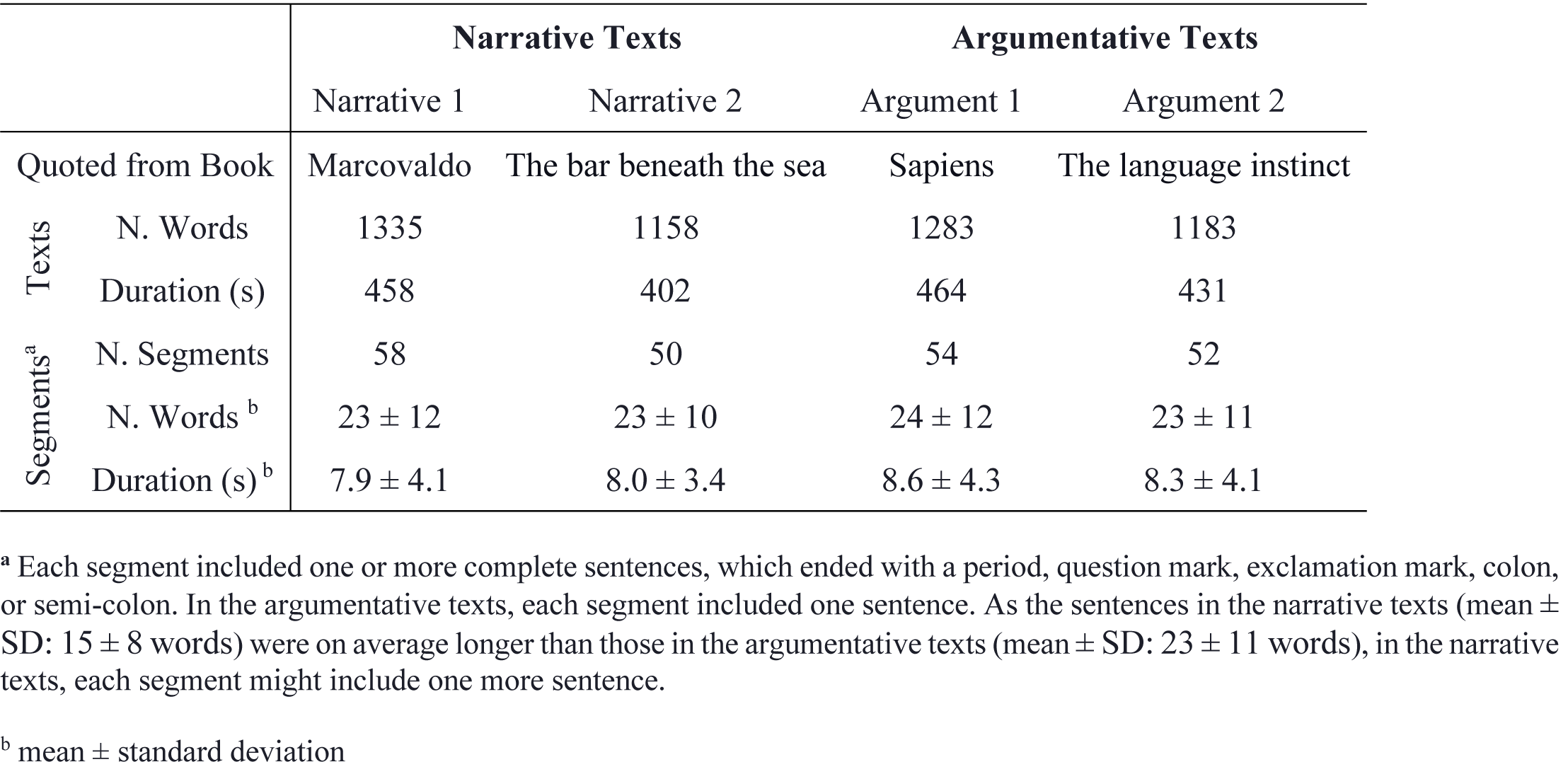
Information on Selected Texts

|  |  | Narrative Texts |  | Argumentative Texts |  |
| --- | --- | --- | --- | --- | --- |
|  |  | Narrative 1 | Narrative 2 | Argument 1 | Argument 2 |
| Quoted from Book |  | Marcovaldo | The bar beneath the sea | Sapiens | The language instinct |
| Texts | N. Words | 1335 | 1158 | 1283 | 1183 |
|  | Duration (s) | 458 | 402 | 464 | 431 |
| Segments<sup>a</sup> | N. Segments | 58 | 50 | 54 | 52 |
|  | N. Words<sup>b</sup> | 23 ± 12 | 23 ± 10 | 24 ± 12 | 23 ± 11 |
|  | Duration (s)<sup>b</sup> | 7.9 ± 4.1 | 8.0 ± 3.4 | 8.6 ± 4.3 | 8.3 ± 4.1 |

<sup>a</sup> Each segment included one or more complete sentences, which ended with a period, question mark, exclamation mark, colon, or semi-colon. In the argumentative texts, each segment included one sentence. As the sentences in the narrative texts (mean ± SD: 15 ± 8 words) were on average longer than those in the argumentative texts (mean ± SD: 23 ± 11 words), in the narrative texts, each segment might include one more sentence.
<sup>b</sup> mean ± standard deviation

## Results

### Behavior rating on stimuli

Table 1 shows the information on the two selected narrative texts and the two selected argumentative texts (see Methods for more detailed information). These texts were divided into segments consisting of complete sentences, each one ending with a period, question mark, exclamation mark, colon, or semi-colon. We sorted the segments according to random order and concatenated them together to generate a sentence-scrambled version for each text. The number of words, duration, number of segments, number of words of each segment, and the duration of each segment were matched between narrative texts and argumentative texts. These measurements were also comparable to those in the previous studies using ISC (Lerner et al., 2011) and ISFC (Simony et al., 2016) methods.

At the stimuli-selection stage, we rated narrative- and argument-relevant features of these texts on a five-point Likert scale (Fig. 1). The questionnaire used to query these features can be found in the supplementary materials. Each text was rated by 20 participants who did not participate in the MRI experiment (see Methods for more detailed information). The results confirmed that the two narrative texts had higher ratings than the two argumentative texts on narrative-related features such as narrativeness (Welch’s t(77.81) = 20.11; P < 0.001), concreteness (Welch’s t(69.93) = 3.39; P = 0.001), scene construction (Welch’s t(52.52) = 9.24; P < 0.001), self-projection (Welch’s t(68.92) = 5.18; P < 0.001), and theory of mind (Welch’s t(77.97) = 3.99; P < 0.001) (Figure 1a). The two argumentative texts received higher ratings than the two narrative texts on argument-related features such as argumentativeness (Welch’s t(78.00) = -10.36, P < 0.001), abstractness (Welch’s t(78.00) = -11.51, P < 0.001), and logical thinking (Welch’s t(77.81) = -11.03, P < 0.001) (Fig. 1b).

**Figure 1.**
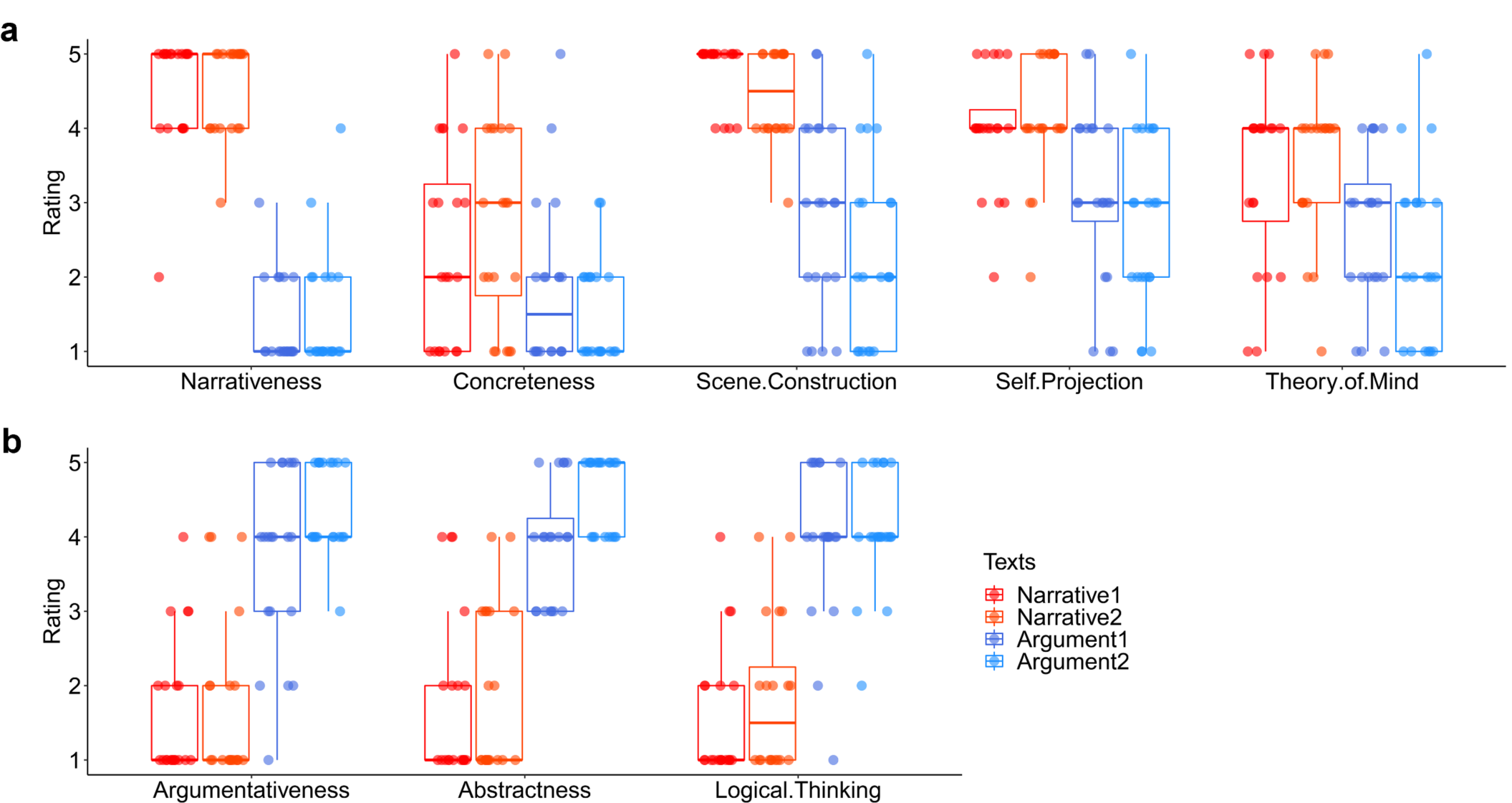
Behavior rating on the four selected texts. The boxplots show the rating scores on the four chosen texts from an independent group of participants who did not participate in the fMRI experiment. Each text was rated by 20 participants. Figure 1a shows the rating scores on the narrative texts were significantly higher than the argumentative texts on the items of narrativeness, concreteness, scene construction, self-projection, and theory of mind. Figure 1b shows the rating scores on argumentative texts were significantly higher than the narrative texts on the items of argumentativeness, abstractness, and logical thinking. The participants who participated in the fMRI experiment did the same rating. The results validated the rating pattern here, as shown in Supplementary Figure 1.

The 16 participants who took part in the fMRI experiment filled in the same rating questionnaire after scanning. The results largely validated the above rating patterns (Supplementary Fig. 1). In the questionnaire, these participants also rated to which degree they understood the texts on a five-point Likert scale. The results showed that they understood the intact texts better than the sentence-scrambled texts: The comprehensibility rating on the intact narrative texts (mean ± SD: 4.69 ± 0.51) was significantly higher than the scrambled narrative texts (mean ± SD: 2.63 ± 0.67) (pair t(15) = 15.17, P < 0.001), and the comprehensibility rating on the argumentative texts (mean ± SD: 4.41 ± 0.74) was significantly higher than the scrambled argumentative texts (mean ± SD: 2.97 ± 0.99) (pair t(15) = 8.46, P < 0.001).

### Narrative, not argumentative texts, evoked time-locked neural activity in the DMN

We first investigated the time-locked regional activity evoked by narrative and argumentative thought by comparing the ISC in the intact-text conditions when the participants could construct coherent thoughts to the ISC in the scrambled-sentence conditions when participants could only process the literal meaning of each sentence (Fig. 2). To recognize which brain systems are engaged in narrative and argumentative thought, we calculated the percentage of significant brain areas (i.e., the number of vertexes) that fall into each pre-identified brain system. The distribution of each brain system was identified based on a study applying clustering analysis on the interregional connectivity pattern (Thomas Yeo et al., 2011) (Supplementary Fig. 2; see Methods for details).

**Figure 2.**
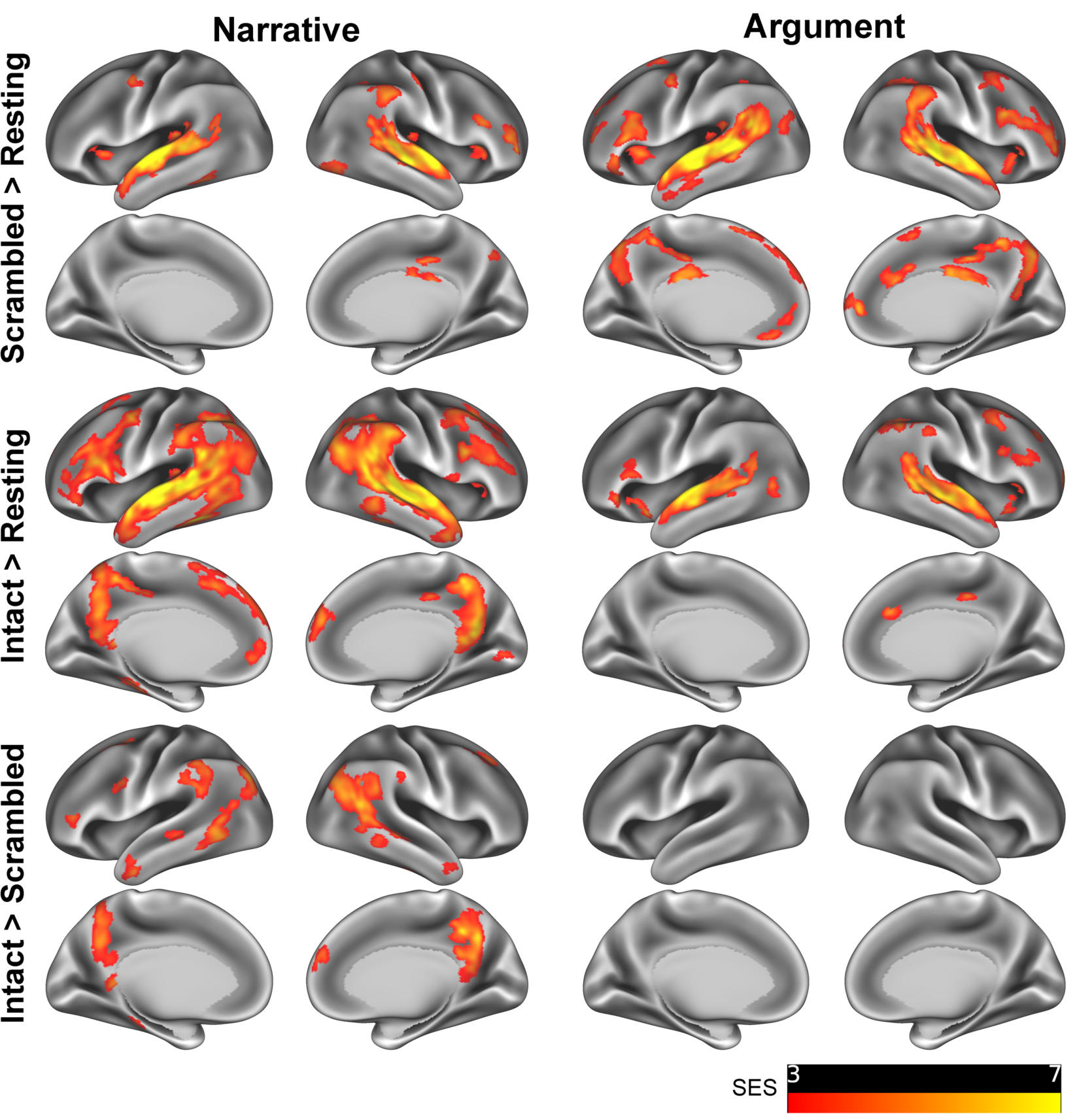
ISC revealed the narrative-induced activity in DMN but not the argumentative one. ISC contrast maps illustrate the significant areas of each contrast in the narrative (left) and the argumentative (right) conditions (P < 0.05, FDR corrected, Area > 200 mm^2^). The first row shows the results in the contrast between the scrambled-sentence conditions and the resting state. For both narrative and argumentative conditions, mainly the auditory system, the language system, and the domain-general system are involved. The second row shows the results of the contrast between the intact-text conditions and the resting state. While the neural distribution in the intact-narrative condition extended to other brain systems like the DMN, the neural distribution in the intact-argumentative condition was confined to the areas in the scrambled-argumentative condition. The third row shows the results of the direct contrast between the intact-text condition and the scrambled-sentence condition. Areas in the default mode, language, control, and attention systems were more engaged in the intact narratives. We did not find any significant areas in this contrast for the argumentative condition. SES: standard effect size.

As a sanity check, we examined the contrast between the scrambled-sentence condition and the resting-state condition. We predicted that sentence-scrambled texts should mainly synchronize the auditory, language, and domain-general process across participants. The results confirmed this prediction by showing that, independently of text type (narrative or argumentative), about 90% of significant vertexes fell into the four brain systems relating to auditory, language, control, and attention (P < 0.05, FDR corrected, area > 200 mm^2^; Fig. 2, first row; Supplementary Fig. 2b, first row).

We moved on to investigate the neural correlates of narrative and argumentative thought by detecting the regions that show additional or higher synchronization in the intact-text condition compared to the scrambled-sentence condition (P < 0.05, FDR corrected, area > 200 mm^2^; Fig. 2, second and third row; Supplementary Fig. 2b, second and third row). The results contrasting intact-narrative condition to the resting-state condition showed a much wider distribution of brain areas than the results contrasting scrambled-sentence condition to the resting-state condition. Note that, in the intact-narrative condition, 18% of significant regions fell into the DMN, whereas in the scrambled-narrative condition, this portion was less than 1%. Directly contrasting the intact-narrative condition to the scrambled-narrative condition revealed about 90% of significant regions fell in four brain systems: the default mode, language, control, and attention, of which 37% were in the DMN. Specifically, the significant regions in the DMN included the angular gyrus (AG), the area comprising the precuneus, the posterior cingulate cortex (PCC), and the ventral retrosplenial complex (RSC), and the middle portion of the left peri-hippocampal area. Intriguingly, contrasting intact-argumentative condition to the resting-state condition only showed brain areas confined within the brain areas that emerge when contrasting the scrambled-sentence condition to the resting-state condition. Directly contrasting the intact-argumentative condition to the scrambled-argumentative condition did not reveal any additional brain areas, even at a lower threshold (P < 0.001, uncorrected).

We also contrasted the ISC result of narrative thought to the argumentative one, i.e., (Intact Narrative - Scrambled Narrative) > (Intact Argument - Scrambled Argument) (P < 0.05, FDR corrected, area > 200 mm^2^; Supplementary Fig 3). The significant brain areas coincided with the results of the narrative thought: Over 90% of the significant regions fell into the default mode, language, control, and attention systems, of which 25% were in the DMN. The opposite contrast did not reveal any region more involved in the argumentative thought than the narrative one, even at a lower threshold (P < 0.001, uncorrected).

**Figure 3.**
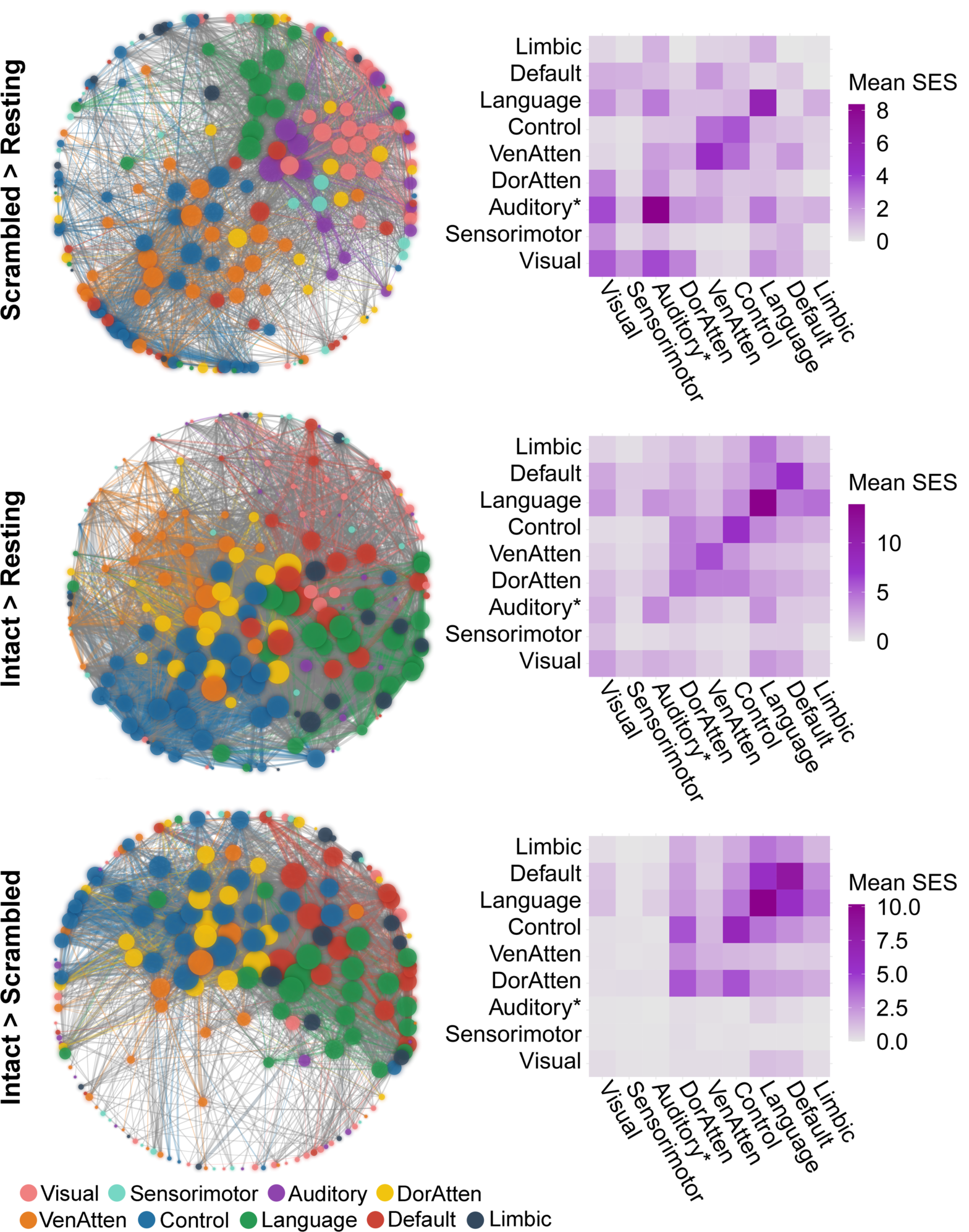
Network reconfiguration for narrative comprehension. The figure illustrates the significant ISFC difference (P < 0.05, FWE corrected) between the scrambled-narrative condition and the resting state (the first row), the intact-narrative condition and the resting state (the second row), and the intact- and scrambled-narrative condition (the third row). The left column shows the network layout, where the nodes represent the brain areas, and the edges represent the significant interregional ISFC differences between conditions. This layout was generated using the force-directed graph drawing algorithm: strongly connected nodes cluster together, and weakly connected nodes are pushed apart. The size of the nodes denotes the node degree of each brain area. The color of the nodes denotes to which brain system they belong. The width of the edges denotes the standardized effect size (SES). Intra-system edges are in the color of that network; inter-system edges are in gray. The right column shows the distribution of all the significant edges of each contrast within or between brain systems. Each cell indicates the mean SES of each contrast, i.e., the ratio between the sum of the SES and the number of the edges in the fully connected situation. “Auditory*” denotes the network including not only the auditory cortex but also the ventral somatosensory and motor brain areas corresponding to the body parts above the neck. VenAtten = Ventral Attention; DorAtten = Dorsal Attention.

To validate the above results and to evaluate the across-stimuli consistency, we repeated the analysis on each of the two narrative texts (Supplementary Fig. 4) and the two argumentative texts (Supplementary Fig. 5) (P < 0.05, FDR corrected, area > 200 mm^2^). The results showed an overall consistency between the two texts of the same type despite the considerable difference in content and writing style. For the two narrative texts, contrasting the intact-text condition to the scrambled-sentence condition revealed significant brain areas that mostly overlapped in the DMN, i.e., the precuneus and the posterior angular gyrus. For the two argumentative texts, the same contrast did not reveal any significant brain areas.

**Figure 4.**
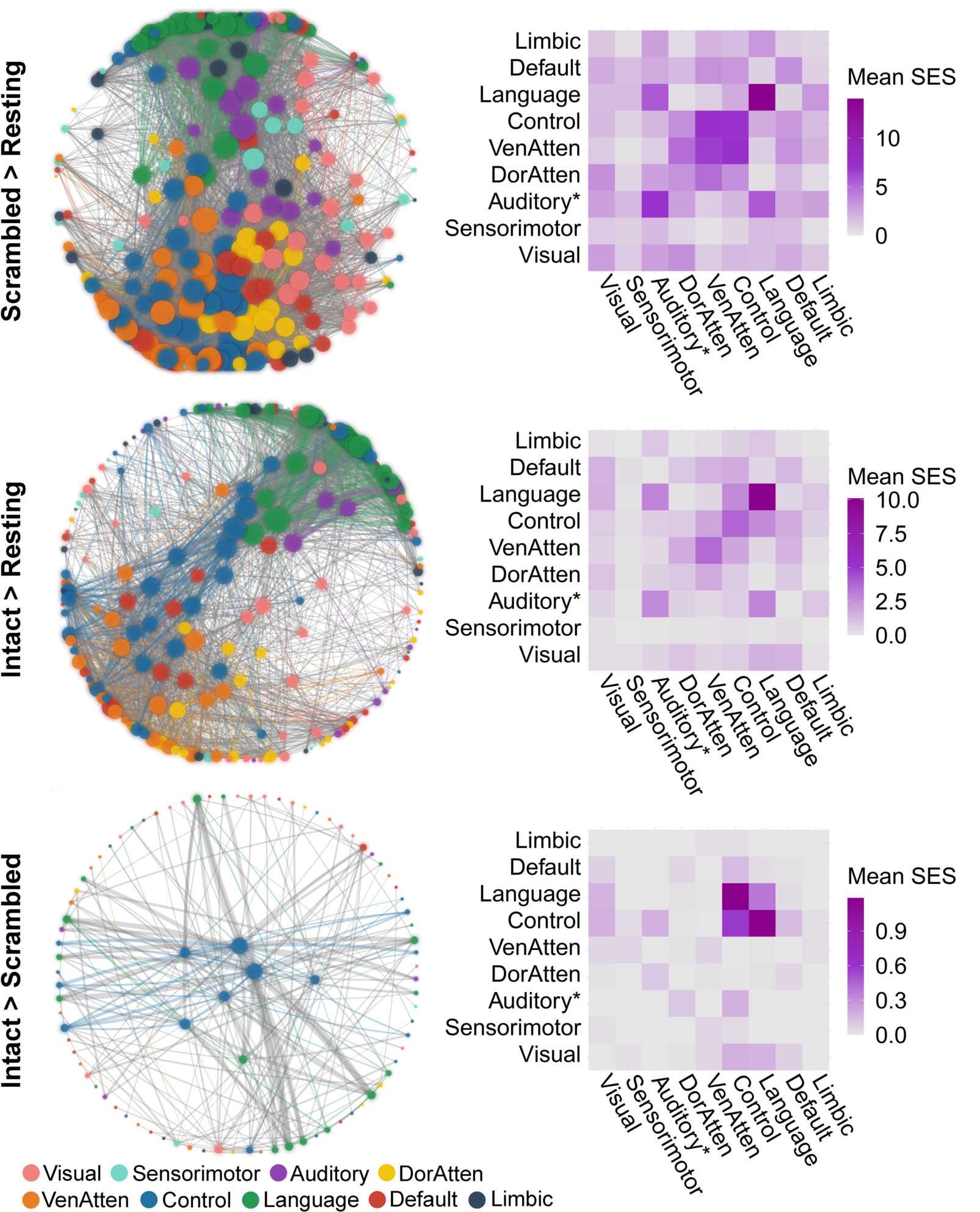
Network reconfiguration for argumentative comprehension. The figure illustrates the significant ISFC difference (P < 0.05, FWE corrected) between the scrambled-argumentative condition and the resting state (the first row), the intact-argumentative condition and the resting state (the second row), and the intact- and scrambled-argumentative conditions (the third row). The left column shows the network layout, where the nodes represent the brain areas, and the edges represent the significant inter-regional ISFC differences between conditions. This layout was generated using the force-directed graph drawing algorithm: strongly connected nodes cluster together, and weakly connected nodes are pushed apart. The size of the nodes denotes the node degree of each brain area. The color of the nodes denotes to which brain system they belong. The width of the edges denotes the standardized effect size (SES). Intra-system edges are in the color of that network; inter-system edges are in gray. The right column shows the distribution of all the significant edges of each contrast within or between brain systems. Each cell indicates the mean SES of each contrast, i.e., the ratio between the sum of the SES and the number of the edges in the fully connected situation. “Auditory*” denotes the network including not only the auditory cortex but also the ventral somatosensory and motor brain areas corresponding to the body parts above the neck. VenAtten = Ventral Attention; DorAtten = Dorsal Attention.

**Figure 5.**
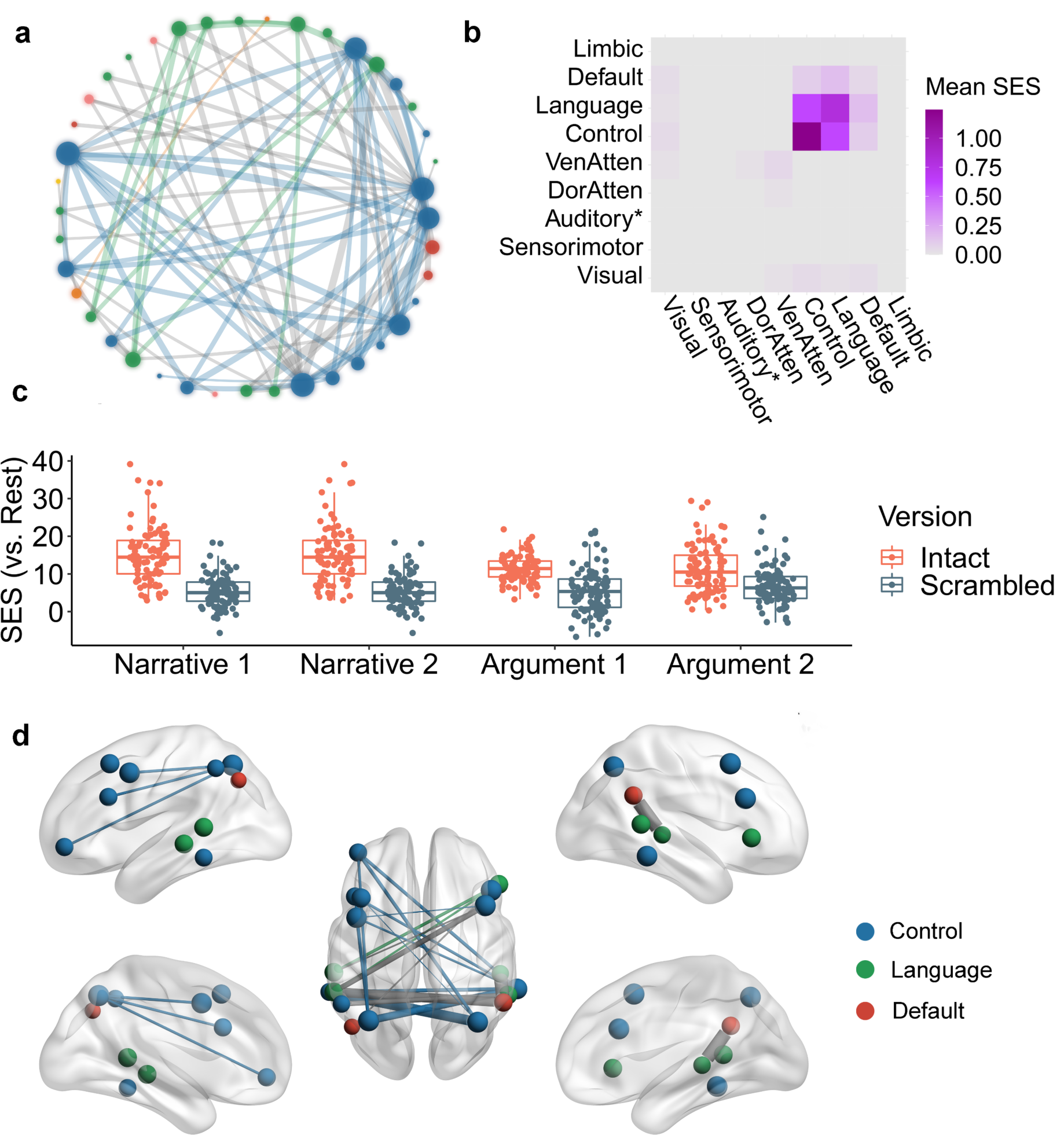
The shared network for both narrative and argumentative thought. Figure a illustrates the network layout of the shared brain network for narrative and argumentative thought using the force-directed graph drawing algorithm. It consisted of 88 edges. The color legend is the same as the one in Figure 3 and Figure 4. Figure b illustrates the distribution of 88 edges within and between brain systems, where each cell indicates the mean standardized effect size (SES) in the contrast between the narrative and the scrambled conditions, i.e., the ratio between the sum of the SES and the number of the edges in the fully connected situation. Figure c illustrates the SES of the 88 edges of all the conditions in contrast to the resting state. Figure d illustrates the top 20 edges with the largest SES in the contrast between narrative and scrambled conditions. In Figure a and Figure d, the size of the nodes denotes the node degree of each brain area in the whole graph comprising 88 edges. The color of the nodes denotes to which brain system they belong. The width of the edges denotes the SES. Intra-system edges are in the color of that network; inter-system edges are in gray.

The above ISC analysis verifies the previous findings that the DMN engages in narrative thought (Ferstl et al., 2008; Lerner et al., 2011), but fails to reveal the neural basis for the argumentative one. It seems that the DMN does not serve as the general machinery for long-timescale information integration, supporting both modes of thought.

### Network reconfiguration for narrative and argumentative thought

It is worth noting that the ISC analysis investigates the stimulus-induced neural activity region by region in isolation. Constructing a coherent thought throughout a relatively long text might rely on the reconfiguration of brain networks already active during sentence-level processing, without necessarily recruiting additional brain regions. As the ISFC measures the purely stimulus-induced functional coupling between discrete regions (Simony et al., 2016), it can reflect the brain network reconfigurations across different task states. The current analysis aimed to investigate the network reconfiguration for narrative and argumentative thought by comparing the ISFC in the intact-text conditions to those in the scrambled-sentence conditions (Fig. 3 and Fig. 4).

We implemented the ISFC analysis based on a whole-brain parcellation atlas comprising 200 brain regions (Schaefer et al., 2018). The atlas also provides information about which brain system each of the 200 brain areas belong to. Fig. 3 and Fig. 4 illustrate the network reconfiguration in the narrative conditions and the argumentative conditions, respectively. The left panel in both figures shows the network layout of all the significant ISFC differences between conditions (P < 0.05, FWE corrected) using the force-directed graph drawing algorithm (Fruchterman and Reingold, 1991), where strongly connected nodes cluster together, and weakly connected nodes are pushed apart. The nodes represent brain areas of each brain system, where the size of nodes denotes the node degree, i.e., the sum of edges that connect to the nodes. The edges represent the significant interregional ISFC difference, where the width of edges denotes the standardized effect size of the contrast (SES). The right panel in both figures summarizes the edge distribution within and between brain systems. Each cell denotes the mean SES, i.e., the ratio between the sum of all the significant edges and the number of all the possible edges in the fully connected situation.

For narrative conditions, the ISFC results were mostly in line with the ISC results. Scrambled-narrative texts, in contrast to the resting state, synchronized the neural activity mainly in the brain systems relating to auditory, language, control, and ventral attention (Fig. 3, the first row). Intact-narrative texts, in contrast to the resting state, extended the synchronization to the DMN (Fig. 3, the second row). A direct comparison between the intact-narrative condition and the scrambled-narrative condition was implemented by detecting the edges that simultaneously met the criteria (1) Intact Narrative > Scrambled Narrative (P < 0.05, FWE corrected) and (2) Intact Narrative > Resting State (P < 0.05, FWE corrected). The significant edges mainly fell into the brain systems relating to the default mode, language, control, and dorsal attention. (Fig. 3, the third row). Supplementary Fig. 6a illustrates the top 20 edges with the biggest SES within the DMN. These critical functional couplings covered all the core regions in the DMN, i.e., the AG (Brodmann area 39), the dorsal lateral prefrontal cortex (8Ad area) (Petrides, 1999), the anterior medial prefrontal cortex, the PCC, the ventral RSC, and the parahippocampal area. The result confirmed previous findings that areas in the DMN are synchronized as a network to support the narrative thought (Simony et al., 2016).

**Figure 6.**
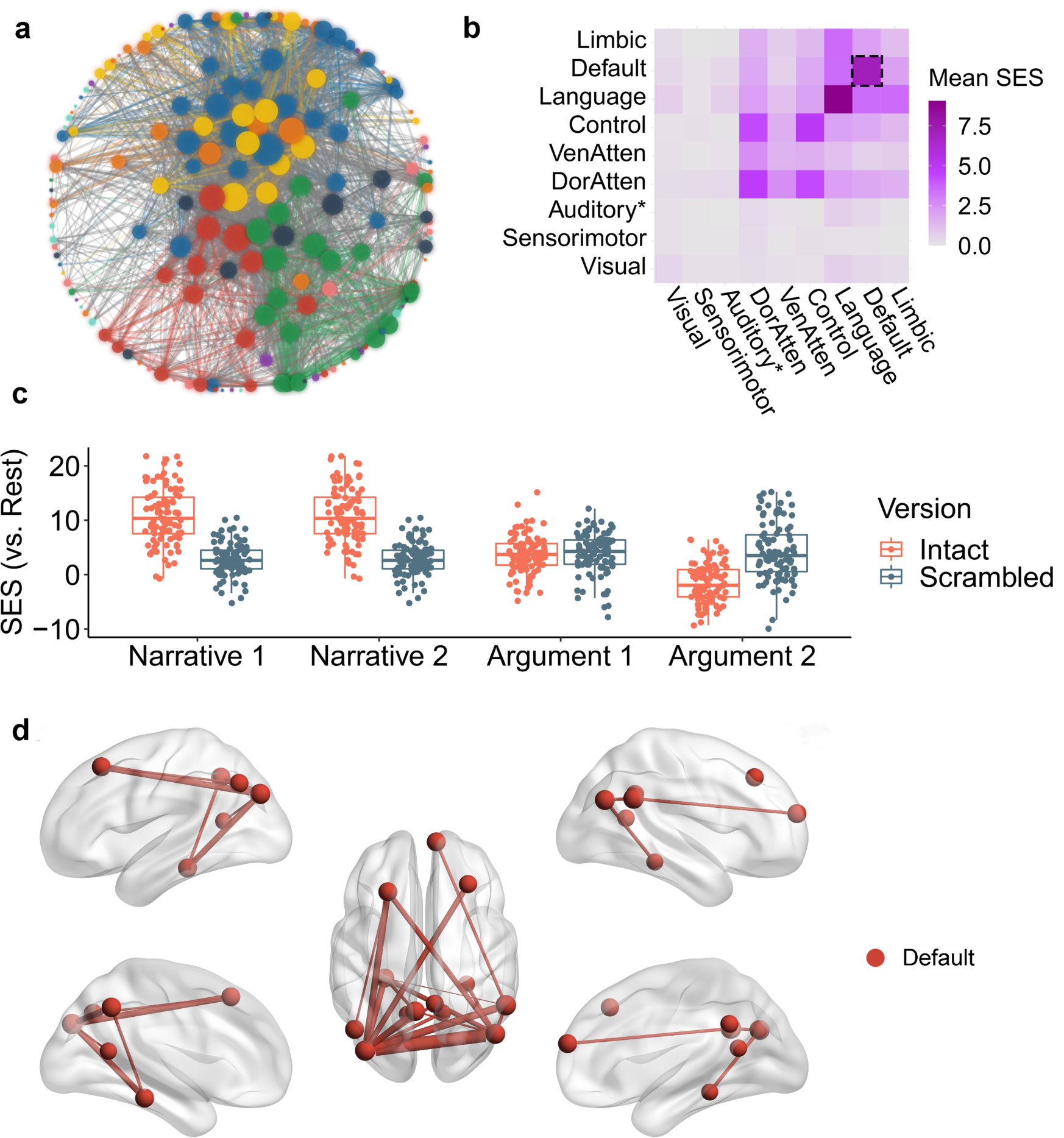
The brain network specific to narrative thought. Figure a illustrates the network layout of the brain network more sensitive to narrative thought using the force-directed graph drawing algorithm. It consisted of 2348 edges. The color legend is the same as the one in Figure 3 and Figure 4. Figure b illustrates the distribution of the 2348 edges within and between brain systems, where each cell indicates the mean standardized effect size (SES) in the “(intact-narrative – scrambled narrative) > (intact argumentative – scrambled argumentative)” contrast, i.e., the ratio between the sum of the SES and the number of the edges in the fully connected situation. There were 96 edges in DMN, which are highlighted using the dotted lines. Figure c illustrates the SES of the 96 edges in the DMN of all the conditions in contrast to the resting state. Figure d illustrates the top 20 edges within the 96 edges in the DMN with the largest SES in the “(intact-narrative – scrambled-narrative) > (intact-argumentative – scrambled-argumentative)” contrast. In Figure a and Figure d, the size of the nodes denotes the node degree of each brain area in the whole graph comprising 2348 edges. The color of the nodes denotes to which brain system they belong. The width of the edges denotes the SES. Intra-system edges are in the color of that network; inter-system edges are in gray.

For argumentative conditions, scrambled-argumentative texts, in contrast to the resting state, also synchronized the neural activity mainly in the brain systems relating to auditory, language, control, and ventral attention (Fig. 4, the first row). Intact-argumentative texts seemed not to involve additional brain systems. Most of the significant edges were within the language system (Fig. 4, the second row). A direct comparison between the intact-argumentative condition and the scrambled-argumentative condition was conducted by detecting the edges that simultaneously met the criteria (1) Intact Argument > Scrambled Argument (P < 0.05, FWE corrected) and (2) Intact Argument > Resting State (P < 0.05, FWE corrected). The significant edges were mostly within the control system or connected the control and the language systems. Not even a single significant edge fell into the DMN. Supplementary Fig. 6b illustrates the top 20 edges with the biggest SES in all the brain systems. All these critical functional couplings were between the control system and the language systems. More specifically, they were the one-to-many connections from the bilateral anterior bank of the intraparietal sulcus (IPS) in the control system to multiple perisylvian areas in the language system including the orbital frontal cortex (Brodmann area 47), the dorsal lateral part of the temporal pole, the whole length of superior temporal gyrus/sulcus (STG/STS, Brodmann area 22), and the temporoparietal junction (TPJ).

We also validated the above results and evaluated the inter-stimuli consistency within the same text type by repeating the analysis on each of the two narrative texts (Supplementary Fig. 7) and each of the two argumentative texts (Supplementary Fig. 8). The results indicated a substantial level of consistency between the different texts of the same type.

**Figure 7.**
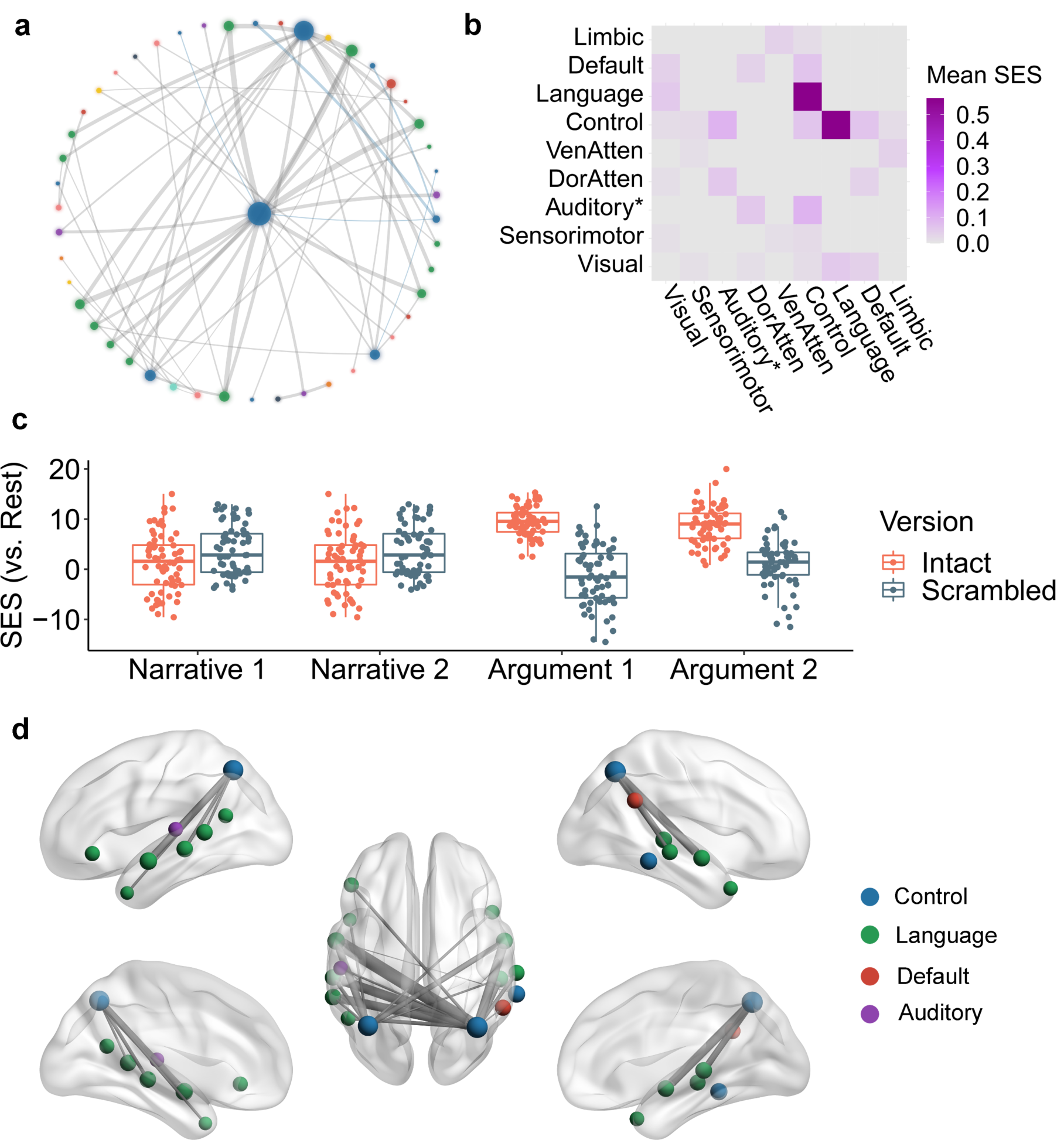
The brain network specific to argumentative thought. Figure a illustrates the network layout of the brain network specific to argumentative thought using the force-directed graph drawing algorithm. It consisted of 64 edges. The color legend is the same as the one in Figure 3 and Figure 4. Figure b illustrates the distribution of 64 edges within and between brain systems, where each cell indicates the mean standardized effect size (SES) in the “(intact argumentative – scrambled argumentative) > (intact-narrative – scrambled narrative)” contrast, i.e., the ratio between the sum of the SES and the number of the edges in the fully connected situation. Figure c illustrates the SES of the 64 edges of all the conditions in contrast to the resting state. Figure d illustrates the top 20 edges with the largest SES in the “(intact argumentative – scrambled argumentative) > (intact-narrative – scrambled narrative)” contrast. In Figure a and Figure d, the size of the nodes denotes the node degree of each brain area in the whole graph comprising 64 edges. The color of the nodes denotes to which brain system they belong. The width of the edges denotes the SES. Intra-system edges are in the color of that network; inter-system edges are in gray.

### Commonalities and differences between narrative and argumentative networks

Next, we disentangled the brain network shared by both narrative and argumentative thought from the brain network specific to narrative or argumentative thought. The shared brain network for both narrative and argumentative thought was defined as the functional couplings which met the following criteria simultaneously: (1) Intact Narrative > Resting State (P < 0.05, FWE corrected); (2) Intact Narrative > Scrambled Narrative (P < 0.05, FWE corrected); (3) Intact Argument > Resting State (P < 0.05, FWE corrected); (4) Intact Argument > Scrambled Argument (P < 0.05, FWE corrected). We found 88 edges that meet these criteria (Figure 5a). Most of the functional couplings were in the control system; the others were mainly within the language system or between the language and control system (Figure 5b). Figure 5c illustrates the SES of the 88 edges of the contrast between each condition and the resting state. The SES in the intact condition was greater than the one in the scrambled condition for all the four texts regardless they were narrative or argumentative. Figure 5d illustrates the top 20 edges with the largest averaged SES in the contrasts between intact-narrative condition and scrambled-narrative condition and between intact-argumentative condition and scrambled-argumentative condition. Most edges linked areas within the control system. They connected the anterior bank of the IPS to multiple lateral prefrontal regions and the temporooccipital area at the temporal entrance.

The brain network more sensitive to narrative thought was defined as the functional coupling which met the following criteria: (1) (Intact Narrative – Scrambled Narrative) > (Intact Argument – Scrambled Argument) (P < 0.05, FWE correction); (2) Intact Narrative > Scrambled Narrative (P < 0.05, FWE correction); (3) Intact Narrative > Resting State (P < 0.05, FWE correction). We found 2348 edges that met these criteria (Figure 6a). These edges mainly related to the language, default mode, control, and dorsal attention systems (Figure 6b). There were 96 edges in the DMN. Figure 6c illustrates the SES of these 96 edges of the contrast between each condition and the resting state. The SES in the intact-narrative conditions was greater than the one in the scrambled-narrative conditions. However, the SES in the intact-argumentative conditions was not greater than the one in the scrambled-argumentative conditions. Figure 6d illustrates the top 20 edges in the DMN with the largest SES in the “(Intact Narrative – Scrambled Narrative) > (Intact Argument – Scrambled Argument)” contrast. These edges covered all the core regions in the DMN, including the AG (Brodmann area 39), the dorsal lateral prefrontal cortex (8Ad area) (Petrides, 1999), the anterior medial prefrontal cortex, the PCC, the ventral RSC, and the parahippocampal area.

The brain network specific to argumentative thought was defined as the functional coupling which met the following criteria: (1) (Intact Argument - Scrambled Argument) > (Intact Narrative – Scrambled Narrative) (P < 0.05, FWE correction); (2) Intact Argument > Scrambled Argument (P < 0.05, FWE correction); (3) Intact Argument > Resting State (P < 0.05, FWE correction). We found 64 edges that met these criteria (Figure 7a). These edges mainly connected the control and the language systems (Figure 7b). Figure 7c illustrates the SES of these 64 edges of the contrast between each condition and the resting state. The SES in the intact-argumentative conditions was greater than the one in the scrambled-argumentative conditions. However, the SES in the intact-narrative conditions was not greater than the one in the scrambled-narrative conditions. Figure 7d illustrates the top 20 edges in the whole brain with the largest SES in the “(Intact Argument – Scrambled Argument) > (Intact Narrative – Scrambled Narrative)” contrast. Most of the edges connected the control system and the language systems, more specifically, the one-to-many connections between the bilateral anterior bank of the IPS in the control system and multiple perisylvian areas in the language system, including the orbital frontal cortex (Brodmann area 47), the dorsal lateral part of the temporal pole, the whole length of STG/STS (Brodmann area 22), and the TPJ.

## Discussion

To investigate the neural basis of the narrative and argumentative thought, we compared the stimuli-evoked regional neural activity and functional coupling when participants were listening to narrative and argumentative texts to those when participants were listening to sentence-scrambled text. We found that the sentence-scrambled texts, whether they were narrative or argumentative, induced regional neural activity and functional coupling mainly in the brain systems relating to auditory, language, attention, and control (first row in Fig. 2, Fig. 3, and Fig. 4). While the intact-narrative condition additionally involved the DMN, the intact-argumentative condition did not extend to other brain systems (second row in Fig. 2, Fig. 3, and Fig. 4). Directly contrasted to the scrambled-sentence conditions, both intact-narrative condition and intact-argumentative condition enhanced functional coupling mainly in the frontoparietal control system (Fig. 5). The intact-narrative condition in contrast to the scrambled-narrative condition also induced widely distributed neural activity and functional coupling that implicated the core regions in the DMN (third row in Fig. 2 and Fig. 3; Fig. 6). However, we failed to find any neural activity or functional coupling in the DMN when contrasting the intact-argumentative condition to the scrambled-argumentative condition (third row in Fig. 2 and Fig. 4; Fig. 6c). Instead, we found functional couplings between the anterior bank of the IPS in the control system and multiple perisylvian areas in the language system (Fig. 7). These one-to-many connections were not significant in the contrast between the intact-narrative condition and the scrambled-narrative condition (Fig. 7c). We also validated our results by implementing the same analyses on each of the two selected texts of the same types. The pattern of the result was consistent with the one pooling two texts together (Supplementary Fig. 4, Fig. 5, Fig. 7, Fig. 8).

The results revealed the commonalities and differences between neural bases underlying narrative and argumentative thought, which seems to support both content-independent and content-dependent hypotheses (Jacoby and Fedorenko, 2018). The content-independent hypothesis predicts that the narrative and the argumentative thoughts should share the same neural basis because the coherence of both modes of thought relies on iteratively accumulating and updating information over a long timescale. Instead of the DMN, we found the shared neural basis for both narrative and argumentative thought in the frontoparietal control system. The frontoparietal control system, together with the attention-relevant regions in cingulo-opercular areas, is usually referred to as the “multiple demand system” (Duncan, 2010), which is named after its broad engagement in a wide variety of demanding tasks (Fedorenko et al., 2013). However, unlike the sustained activity in the attention-relevant brain area, the frontoparietal control system rapidly adjusts its activity profile (MacDonald et al., 2000) and global functional connectivity pattern (Cole et al., 2013) to adapt to the task context. Our results suggest that both modes of thought may rely on the frontoparietal control system as a general working memory system to iteratively accumulating and updating information over long temporal windows.

The content-dependent hypothesis predicts that the neural bases underlying narrative and argumentative thought are irreducible to each other as these two modes of thought differ in their core cognitive components. As mentioned in the Introduction, the narrative thought relies on constructing and updating a “situation model” about the state of affairs to understand the temporal causality of the events and the intention of the characters (Zwaan and Radvansky, 1998). The argumentative thought, instead, relies on “informal logic” processing, which includes identification and evaluation of the logic structures that are embedded in the natural language discourse (Blair, 2015). The findings that the DMN was specific for narrative thought, and the cooperation between the control and the language systems via the IPS was specific for argumentative thought, may support this hypothesis. The functionality of situation model construction coincides with the role of the DMN in scene construction (Hassabis and Maguire, 2007; Spreng et al., 2009), self-projection (Buckner and Carroll, 2007; Spreng et al., 2009), prospection (Schacter et al., 2007; Spreng et al., 2009), and theory of mind (Lin et al., 2018; Spreng et al., 2009). The IPS, together with other brain areas in the frontoparietal control systems, is considered as the neural basis of fluid intelligence (Bishop et al., 2008). Thus, the coordination and cooperation between the frontoparietal control system and the language system, which is mediated by the IPS, might be critical to identify and evaluate the informal logic in the natural language discourse.

How to reconcile these two seemingly opposing hypotheses? A likely possibility is that the brain function is simultaneously featured by two factors: the temporal receptive window (TRW) for information processing and the information types. Take the frontoparietal control system and the default mode system as an example. On the one hand, according to the hierarchical process memory framework (Hasson et al., 2015), the TRW of a brain system is defined by its position in the cortical hierarchy. In terms of connectivity pattern, the frontoparietal control system and the default mode system are at the medial and top level of the cortical hierarchy, respectively (Margulies et al., 2016; Sepulcre et al., 2012). They thus can process longer-timescale information (e.g., the “train of thought”) than the sensorimotor cortices at the low level of the cortical hierarchy. On the other hand, the information type processed by a brain system is defined by its wiring patterns to the other functionally specialized brain modules. The frontoparietal control system, which has widely distributed connections to the other brain systems (Power et al., 2011), can serve as the general machinery to integrate long-timescale information of all kinds. The default mode system, which has strong connections mainly to the medial temporal lobe, is more likely an extension to the episodic memory system (Buckner et al., 2008), which is sensitive to the narrative information. Given the default mode system is at an even higher level of in the cortical hierarchy than the frontoparietal control system (Margulies et al., 2016; Sepulcre et al., 2012), the default mode system could have the capacity to process longer narrative information than the domain-general information which is processed by the frontoparietal control system. If this is true, it might be the reason to explain why narratives tend to be more accessible and memorable than the other genres (Graesser et al., 1980; Zabrucky and Moore, 1999).

Our study also indicates the importance of treating the brain as a network and illustrates how diverse mental activities arise from network reconfiguration. There are two general mechanisms at play (Park and Friston, 2013). One mechanism is through local integration. The brain was organized into functionally specialized modular structures, where the areas within the module are densely connected (Thomas Yeo et al., 2011). Each module can be selectively recruited as a functional unite according to task requirements by enhancing its within-module functional couplings. For example, compared to the scrambled-text condition, the intact-narrative condition selectively involved the default mode system (Figure 3) by inducing the functional couplings among all the core regions with the DMN (Supplementary Fig 6a; Fig. 6). Another mechanism is through global integration, which means these recruited modules are coordinated by inter-module connections, aiming to achieve more complicated tasks. Unlike the dense intra-module connections, these inter-module connections are looser, and are usually mediated by a small number of brain areas, termed “connectors.” A prominent example is the neural basis of argumentative thought. In the scrambled-argumentative condition, the language and control systems were already involved but segregated (Figure 4). The intact-argumentative condition did not recruit additional brain systems. Instead, it promoted the cooperation between the control and language systems, and this cooperation is achieved strictly through the IPS, as the connector (Supplementary Fig 6b; Fig 7). The global integration of the local integration strategy guarantees the efficiency and flexibility of brain function, where the functionally specialized brain modules can be combined and coordinated to adapt diverse task context.

To conclude, our study revealed the commonalities and differences in brain network reconfiguration for the narrative and the argumentative thought. While both modes of thought rely on the frontoparietal control system, the narrative thought specifically implicates the DMN, and the argumentative thought specifically requires the cooperation between the control and the language systems, mediated by the IPS. These results provide insights into how the brain generates diverse mental activity through global and local brain network integration.

## Methods

### Participants

Twenty native Italian speakers who had no history of neurological or psychiatric disorders participated in the fMRI experiment. They were paid as compensation for their time. Following the experimental protocol approved by the local ethical committee at the University of Trento, all participants provided informed written consent before the start of the experiment. Data from four participants were discarded: One participant performed badly in the post-scanning questionnaire concerning the content of the narrative and argumentative texts used in the experiment (his/her accuracy was outside 1.5 times the interquartile range below the lower quartile across participants (Supplementary Fig. 9)). Three participants were excluded due to excessive head motion; In two cases, the mean frame displacement index (Power et al., 2014) of functional images was outside 1.5 times the interquartile range above the upper quartile across participants (Supplementary Fig. 8), and one’s structure image was so blurry that failed to be segmented. The remaining 16 participants (9 females; age range: 21 to 31, mean age: 24) were all educated (university students or above) and right-handed (laterality quotient range: +40 to +100; mean: +90) (Oldfield, 1971). This sample size was in line with the studies employing ISC (Lerner et al., 2011) and ISFC (Simony et al., 2016) methods (11 and 18 participants, respectively).

### Stimuli

This study employed a two (narrative vs. argumentative text) by two (intact vs. sentence-scrambled version) design. We generated two stimuli for each of these four conditions following the procedure below. First, we searched for narrative and argumentative texts that met the following criteria: (1) Written in modern Italian. (2) Easy to understand. All the texts come from best-sellers for non-expert readers. (3) Typical. The narrative text includes a story with the typical elements of the story grammar (Rumelhart, 1975): settings, characters, the initialing event, conflicts/goals, actions, and resolutions. The argumentative text includes the interlinked premiss-illative-conclusion argumentative structure (Hitchcock, 2007), with an overall conclusion at the beginning or end of the text. (4) Self-content. The narrative text should be a complete and independent story; the argumentative text should support a conclusion based on the points independent from the previous chapters. (5) Text length between 1000 to 1300 words. We posited that a comfortable speed range for an Italian audiobook is between 165 and 170 words per minute, which is slightly slower than the average speed of the *Radiotelevisione Italiana* (192.46 words per minute) (Rodero, 2012). This criterion ensures the duration of the selected texts is relatively the same, which is about 6 to 8 minutes, comparable to the 7-minute one used in the studies employing ISC (Lerner et al., 2011) and ISFC (Simony et al., 2016) methods. In the end, we preselected seven such texts - three narrative and four argumentative.

Then, we recruited 35 native Italian speakers (who did not participate in the fMRI experiment; 11 females; age range: 23 to 67, mean age: 32) to rate nine features of these seven texts on a five-point Likert scale. Each participant rated four texts; hence each text was rated by 20 participants. The nine features were difficulty, narrativeness, concreteness, scene construction, Self-projection, theory of mind, argumentativeness, abstractness, and logical thinking (see the questionnaire in the supplementary material). For each text, we also designed two questions on its content before the rating questions to indicate whether the participants had read and comprehended the texts (accuracy rate: 5/8 to 8/8, mean accuracy: 7/8). As all participants provided at least one correct response for each text, we did not exclude any data points. We discarded the texts with high ratings on difficulty (mean rating > 3) and chose two narrative texts and two argumentative texts as our stimuli by maximizing the difference between the ratings of these two text types: the narrative texts had higher ratings on narrative, concreteness, scene construction, and theory of mind; the argumentative texts had higher ratings on argumentativeness, abstractness, and logical thinking (Fig. 1). The two selected narrative texts came from: *The wasp treatment* in the book *Marcovaldo* by Italo Calvino, who tells a story in which the protagonist asks his children to catch wasps and uses them to cure his neighbors’ rheumatism (Narrative 1); *Kulala’s four veils* in the book *The bar beneath the sea* by Stefano Benni, who tells a typical fairy tale (Narrative 2). The two selected argumentative texts were truncated from: *Counting happiness* in the book *Sapiens: a brief history of humankind* by Yuval Noah Harari, who discusses which are the most crucial factors leading to happiness (Argument 1); *An instinct to acquire an art* in the book *The language instinct: how the mind creates language* by Steven Pinker, who argues the nature of language is an instinct faculty, not a cultural product (Argument 2).

Next, we divided the selected four texts into segments. Each segment included one or more complete sentences, which ended with a period, question mark, exclamation mark, colon, or semi-colon, i.e., we did not divide the sentences into clauses. We matched the extent of fragmentation (i.e., the number of segments and the length of segments) between these two text types (Table 1). In the argumentative text, each segment consisted of only one complete sentence. As the sentences in narrative texts (mean ± SD: 15 ± 8 words) were on average shorter than those in the argumentative texts (mean ± SD: 23 ± 11 words), in the narrative text, each segment might consist of more than one sentence.

After that, the same professional voice actor recorded all the four texts with relatively the same volume, speed, voice, and tone. We cut the audio clips according to the segments that we had divided. The duration of each segment was comparable to the duration of the sentence-scrambled version (7.7 ± 3.5s) used in the studies employing the ISC/ISFC method (Lerner et al., 2011; Simony et al., 2016) and matched between the two text types (Table 1). We sorted these segments according to a random order and concatenated them together to generate a sentence-scrambled version for each text.

Finally, we added the same 10s neutral music before both intact and scrambled versions of the stimuli following previous studies employing ISC (Lerner et al., 2011). The volume of the music tapered to zero before the audio texts started. As an abrupt beginning of the sound may elicit a global arousal response in the brain, a piece of opening music here helped to capture the participants’ attention and to protect the start of the texts from being affected by such an arousal shift. We excluded the neural signal in this music period from the analysis (see fMRI preprocessing).

### Procedures

Participants were told that they would be listening to the intact and the scrambled version of four texts during fMRI scanning. They were instructed to follow and comprehend the texts attentively and were informed that they would be asked to fill in a post-scanning questionnaire on the content of what they have heard. To avoid visual intrusion, we blindfolded the participants and turned off the light in the scanning room.

We presented the audio stimuli using Psychotoolbox-3 (http://psychtoolbox.org/). The sound was delivered through an in-ear headphone. Before the formal scanning, participants were instructed to check the sound in the headphone under the scanning noise. We adjusted the volume for each participant to ensure they could hear the pronunciation clearly but meanwhile did not feel too loud.

The functional scanning included nine runs, one for the eight-minute resting state, four for the sentence-scrambled version of the texts, and four for intact version of the texts. Each task runs presented one single text. To make sure the participants were unable to replay the stimuli in the resting state, we put the resting-state run before all the task runs. To make sure the participants were unable to construct coherent thought in the sentence-scrambled runs based on the intact texts they had already heard, we put the four sentence-scrambled runs before the runs for the intact texts. The order of the four sentence-scrambled runs was randomized across participants. For the same participant, the intact-text runs followed the same order of their corresponding sentence-scrambled runs.

After the scanning, all participants completed a questionnaire on the content of the texts that they had heard during the scanning. We designed two questions for each of the four texts. In the same questionnaire, we also asked the participants to do the ratings that were used in the stimulus-selection stage. They were also asked to rate to which degree they could understand each text on a five-point Likert scale.

### MRI acquisition

MRI data were acquired using a MAGNETOM Prisma 3T MR scanner (Siemens) with a 64-channel head– neck coil at the Centre for Mind/Brain Sciences, University of Trento. Functional images were acquired using the simultaneous multislices echoplanar imaging sequence, the scanning plane was parallel to the bicommissural plane, the phase encoding direction was from anterior to posterior, repetition time (TR) = 1000 ms, echo time (TE) = 28 ms, flip angle (FA) = 59°, field of view (FOV) = 200 mm × 200 mm, matrix size = 100 × 100, 65 axial slices, slices thickness (ST) = 2 mm, gap = 0.2 mm, voxel size = 2 × 2 × (2 + 0.2) mm, multiband factor = 5. Three-dimensional T1-weighted images were acquired using the magnetization-prepared rapid gradient-echo sequence, sagittal plane, TR = 2140 ms, TE = 2.9 ms, inversion time = 950 ms, FA = 12°, FOV = 288 mm × 288 mm, matrix size = 288 × 288, 208 continuous sagittal slices, ST = 1 mm, voxel size = 1 × 1 × 1 mm.

### MRI preprocessing

We performed fMRI preprocessing using *fMRIPrep 1*.*5*.*0* (Esteban et al., 2019), which is based on *Nipype 1*.*2*.*2* (Gorgolewski et al., 2011). Please see the section *MRI preprocessing using fMRIPrep 1*.*50* in the supplementary material, where a boilerplate text directly generated by the fMRIPrep describes the preprocessing steps used in the current study. The first 10s, which was the music period in the task runs, was labeled as the dummy scans; thus, they were excluded from the analysis. As surface-based analysis can significantly improve the spatial localization compared to the traditional volume-based analysis (Coalson et al., 2018), we used the images in the fsaverge5 surface space generated by *fMRIPrep*.

We excluded the non-neuronal signal sources through two steps (Pruim et al., 2015). First, we removed the motion-relevant noise using an Independent Component Analysis based strategy for Automatic Removal of Motion Artifacts (ICA-AROMA) (Pruim et al., 2015). The identified motion-relevant components and the signal components were fit into the same general linear model (GLM) to predict the BOLD signal in each vertex on the brain surface. We estimated the beta coefficients using the *fitglm* function in Matlab 2019a and subtracted the motion-relevant terms from the BOLD signal. In this way, the motion-relevant components were removed “non-aggressively” by preserving the shared variance between the motion-relevant components and the signal components. Then, we further removed the other nuisance variables like the mean timecourses in a conservative mask of the white matter (WM) and the cerebrospinal fluid (CSF), which were extracted by *fMRIPrep*. As a recent study demonstrate that the low-frequency component (0 - 0.01 Hz) makes a significant contribution to the ISC (Kauppi et al., 2010), we did not implement high-pass temporal filtering. Instead, we fitted the quadratic polynomial time trend together with the WM and the CSF timecourse into the same GLM to predict the timecourse resulting from the first step, aiming to remove the signal drift. In the same way, we estimated the beta coefficients and subtracted the WM, the CSF, and the quadratic polynomial terms from the signal.

We implemented the surface smoothing on the resulting images with a full width at half maximum of 8 mm using the mri_surf2surf command in FreeSurfer (http://surfer.nmr.mgh.harvard.edu/). The timecourse in each vertex was then z-normalized across time points to enter the following analyses.

### Brain network identification

We identified the brain systems based on a pre-labeled atlas (Thomas Yeo et al., 2011). The brain systems in this atlas are identified by applying the clustering analysis on the pattern of 1000 young healthy participant’s resting-state functional connectivity (RSFC). The atlas has two versions: one coarse version with seven networks and one fine-resolution version with 17 networks. We chose the fine-resolution version as the start for two reasons. First, the fine-resolution version separates the dorsal somatosensory and motor cortex corresponding to the body parts mainly below the neck from the ventral networks consisting of the auditory cortex and the somatosensory and motor cortex corresponding to the body parts mainly up the neck. This division helps us to differentiate the auditory cortex from most of the somatosensory and motor areas. Second, the fine-resolution version also separates the language network (Fedorenko et al., 2011) and the DMN (Buckner et al., 2008). Previous studies suggest these two networks are dissociated in respective of both activation profile and functional connectivity pattern (Mineroff et al., 2018; Xu et al., 2017, 2016). We merged the Network 14 and Network 17 as the language network, which mainly includes the perisylvian cortex and the 55b area (Fedorenko et al., 2011; Glasser et al., 2016). We merged Network 15 and Network 16 as the DMN, as these two networks largely correspond to the two identified sub-networks of the DMN (Braga and Buckner, 2017). We preserved the labels used in the coarse version of the atlas for the other brain networks. These networks are visual; ventral attention(Corbetta and Shulman, 2002; Fox et al., 2006), which may implicate multiple networks variably referred to as the salience (Seeley et al., 2007) and the cingulo-opercular (Dosenbach et al., 2008); dorsal attention (Corbetta and Shulman, 2002; Fox et al., 2006); frontoparietal control (Dosenbach et al., 2008; Vincent et al., 2008); and limbic. In the end, we obtained an atlas, including nine brain systems (Supplementary Fig. 2a).

### ISC analysis

The ISC was defined as the Pearson’s correlation between the timecourse in the same area of different participants. We calculated the ISC for each vertex each run using a leave-one-participant-out approach. For each participant, we first averaged the timecourses of all the other participants and then correlated this mean timecourse with this participant’s timecourse. The resulting Pearson’s correlation coefficients (one per participant) were Fisher-z transformed using the inverse hyperbolic tangent function before they were averaged as one ISC index. In this way, we obtained one ISC surface map for each of the nine runs.

We contrasted the ISC surface maps between different conditions to obtain a veritable ISC contrast value for each vertex for each contrast. The major contrasts were: (1) Scrambled Narrative Contrast: (Scrambled Narrative 1 + Scrambled Narrative 2) - 2 × Rest; (2) Intact Narrative Contrast: (Intact Narrative 1 + Intact Narrative 2) - 2 × Rest; (3) Narrative Contrast: (Intact Narrative 1 - Scrambled Narrative 1) + (Intact Narrative 2 - Scrambled Narrative 2); (4) Scrambled Argumentative Contrast: (Scrambled Argument 1 + Scrambled Argument 2) - 2 × Rest; (5) Intact Argumentative Contrast: (Intact Argument 1 + Intact Argument 2) - 2 × Rest; (6) Argumentative Contrast: (Intact Argument 1 - Scrambled Argument 1) + (Intact Argument 2 - Scrambled Argument 2); (7) Narrative Specific Contrast: [(Intact Narrative 1 - Scrambled Narrative 1) + (Intact Narrative 2 - Scrambled Narrative 2)] - [(Intact Argument 1 - Scrambled Argument 1) + (Intact Argument 2 - Scrambled Argument 2)]; (8) Argumentative Specific Contrast: [(Intact Argument 1 - Scrambled Argument 1) + (Intact Argument 2 - Scrambled Argument 2)] - [(Intact Narrative 1 - Scrambled Narrative 1) + (Intact Narrative 2 - Scrambled Narrative 2)]. We also implemented similar contrasts using individual narrative texts and individual argumentative texts to validate our results and to evaluate the inter-stimulus consistency. The following ISFC analysis used the same contrasts here.

The statistical likelihood of each contrast was assessed using the subject-wise bootstrapping method, where the exchangeability and independence assumptions are satisfied (G. Chen et al., 2016). In each bootstrapping iteration, the same number of participants were randomly resampled with replacement. The ISC was calculated between the timecourse of one participant and the mean timecourse of the other participants. Here, “the other participants” were those excluding him/herself and the repeated ones of him/herself due to resampling with replacement (Nili et al., 2014). The obtained Pearson’s correlation coefficients (one per participant) were Fisher-Z transformed and averaged. We then contrasted these maps between conditions in the same way as before. This procedure was repeated 5000 times to form a sampling distribution for each contrast. The null distribution of each contrast was generated by subtracting the veritable contrast value from the sampling distribution, and the veritable contrast value was then ranked against the null distribution (Hall and Wilson, 1991). As the null distribution of each contrast of each vertex was symmetrical (the skewness is within ± 1), to provide a quantitative measure of the magnitude across contrasts and vertexes, we calculated the standardized effect size (SES) as (*x* − *μ*)/*σ*, where *x* is the veritable contrast value, *μ* is the mean of the null distribution, and *σ* is the standard deviation of the null distribution (Botta-Dukát, 2018). To obtain a high-resolution P-value given the limited number of resamples, we estimated the right-tail p-value of each contrast by approximating a generalized Pareto distribution to the tail of the null distribution (Knijnenburg et al., 2009). We corrected for multiple comparisons across the entire brain surface using the false-discovery rate (FDR) correction algorithm without the need for the assumption of independence across vertices (Benjamini and Yekutieli, 2001) (P < 0.05).

### ISFC analysis

The ISFC was defined as the Pearson’s correlation between the timecourse in two discrete brain areas from different participants. We defined the brain areas based on the cortical parcellation derived by integrating the local gradient approach, which detects the abrupt transitions in RSFC patterns, and the global similarity approach, which clusters similar global ISFC patterns despite the spatial proximity (Schaefer et al., 2018). Thus, the obtained parcels are locally homogenous and globally match to the brain networks shown above (Thomas Yeo et al., 2011). We chose the template matched to the 17 brain networks and then relabeled them as nine networks of interest. Considering the trade-off between the spatial resolution and the computational load, we chose the cortical parcellation consisting of 200 parcels. The averaged the timecourses across all vertexes in each parcel was used as the timecourse of that parcel.

We calculated the pair-wised, inter-regional ISFC among the 200 parcels for each run using a leave-one-participant-out approach following the previous study (Simony et al., 2016). A 200 by 200 ISFC matrix *C*was obtained for each of the nine runs, where each element in the matrix (e.g., *C*_*ij*_) represents the ISFC strength between each pair of regions (e.g., the ith and the jth brain areas). To calculate the value of *C*_*ij*_, we first averaged the timecourses of all the other participants in the jth area and then correlated this mean timecourse with this participant’s timecourse in the ith area. The resulting Pearson’s correlation coefficients (one per participant) were then Fisher-z transformed, averaged, and assigned to *C*_*ij*_. Note that the *C*_*ji*_ is not necessarily equal to *C*_*ij*_. To make the ISFC measure unidirectional, we symmetrized the ISFC matrix as (C+ *C*^*T*^) / 2, where *C*^*T*^ is the transpose of the matrix C. We contrasted the ISFC matrix between different conditions using the same contrasts in the ISC analysis to obtain a veritable contrast value for each pair of brain areas for each contrast.

The statistical likelihood of each contrast was assessed using a similar subject-wise bootstrapping method shown in the ISC analysis. In each iteration of the bootstrapping, the same number of participants were randomly resampled with replacement. A 200 by 200 ISFC matrix *C* was calculated using the data from this sample, where each element in the matrix (e.g., *C*_*ij*_) represents the ISFC strength between each pair of brain areas (e.g., the ith and the jth brain areas). *C*_*ij*_ was calculated as Pearson’s correlation coefficient between the timecourse in the ith brain area of one participant and the mean timecourse in the jth brain area of the other participants. Here, “the other participants” were those excluding him/herself and the repeated ones of him/herself due to resampling with replacement (Nili et al., 2014). The obtained Pearson’s correlation coefficients (one per participant) were Fisher-Z transformed, averaged, and assigned to *C*_*ij*_. We symmetrized the ISFC Matrix C in the same way as before. We contrasted these final ISFC matrixes between conditions, ending this iteration. This procedure was repeated 5000 times to form a sampling distribution of ISFC contrast value for each pair of brain areas for each contrast. The null distribution of each contrast was generated by subtracting the veritable contrast value from the sampling distribution (Hall and Wilson, 1991). As the null distribution of each contrast of each pair of brain regions was symmetrical (the skewness is within ± 1), to provide a quantitative measure of the magnitude across contrasts and pairs of brain regions, we calculated the SES as(*x* −*μ*)/*σ*, where *x* is the veritable contrast value, *μ* is the mean of the null distribution, and *σ* is the standard deviation of the null distribution (Botta-Dukát, 2018). We controlled the family-wise error (FWE) rate by defining the threshold at the 5% percentile of the null distribution of the maximum across all pairs of brain areas and thresholded the SES matrix by assigning the insignificant brain pairs to zero.

The resulting thresholded SES adjacency matrix in each contrast was modeled as a weighted graph comprising nodes and edges (Fornito et al., 2016); the nodes represent brain areas, and the edges represent the SES of that contrast for the ISFC between each pair of the brain areas. We used the node degree to measure the importance of one brain area in each contrast. The degree of node i was calculated as 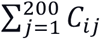, where *C* was the thresholded SES adjacency matrix.

### Visualization

The ISC results were illustrated using the Connectome Workbench 1.3.2 (https://www.humanconnectome.org/software/connectome-workbench). For the visualization purpose, we mapped the significant clusters from the fsaverage5 surface to the fsLR surface using the ADAP_BARY_AREA method. We excluded the clusters that are smaller than 200 mm^2^. The significant clusters were illustrated on an inflate surface against the group-averaged sulcus image of 1096 young adults from the dataset under the Human Connectome Project (https://balsa.wustl.edu/reference/pkXDZ). For the ISFC results, the network layout was generated using the force-directed graph drawing algorithm (Fruchterman and Reingold, 1991) with NodeXL (https://www.smrfoundation.org/nodexl/)(Smith et al., 2010). The brain networks were visualized with the BrainNet Viewer (https://www.nitrc.org/projects/bnv/) (Xia et al., 2013). To localize each node, we used the centroid of the Montreal Neurological Institute coordinates of each brain parcel in the volume version of the same brain parcellation atlas.

## Acknowledgements

We thank Alessia Zampieri for the help in stimuli selection and Anna D’Urso for the help in data collection. This research is supported by the Research Projects of National Interest (PRIN) grant from the Italian Ministry of Education, University and Research (MIUR) to D.C. and O.C. (project number: 2015PCNJ5F_001).

## Author contributions

X.Y. and R.B. conceived the experiment. X.Y. and L.V. analyzed the data. X.Y. wrote the paper with input from L.V., O.C., D.C., and R.B. All authors discussed the results and contributed to the paper.

## Reference

Beach LR, Bissel BL. 2016. Narrative thoughtA New Theory of Mind?: The Thoery of Narrative Thought. Cambridge Scholars Publishing.

Benjamini Y, Yekutieli D. 2001. The control of the false discovery rate in multiple testing under dependency. Ann Stat. doi:10.1214/aos/1013699998

Bishop SJ, Fossella J, Croucher CJ, Duncan J. 2008. COMT val158met genotype affects recruitment of neural mechanisms supporting fluid intelligence. Cereb Cortex 18:2132–2140. doi:10.1093/cercor/bhm240

Blair JA. 2015. What is informal logic?Reflections on Theoretical Issues in Argumentation Theory. Springer. pp. 27–42.

Botta-Dukát Z. 2018. Cautionary note on calculating standardized effect size (SES) in randomization test. Community Ecol. doi:10.1556/168.2018.19.1.8

Braga RM, Buckner RL. 2017. Parallel Interdigitated Distributed Networks within the Individual Estimated by Intrinsic Functional Connectivity. Neuron. doi:10.1016/j.neuron.2017.06.038

Bruner J. 1986. Two modes of thoughtActual Minds, Possible Worlds. Harvard University Press Cambridge MA. pp. 11–43.

Buckner RL, Andrews-Hanna JR, Schacter DL. 2008. The brain’s default network: Anatomy, function, and relevance to disease. Ann N Y Acad Sci 1124:1–38. doi:10.1196/annals.1440.011

Buckner RL, Carroll DC. 2007. Self-projection and the brain. Trends Cogn Sci 11:49–57. doi:10.1016/j.tics.2006.11.004

Chen G, Shin YW, Taylor PA, Glen DR, Reynolds RC, Israel RB, Cox RW. 2016. Untangling the relatedness among correlations, part I: Nonparametric approaches to inter-subject correlation analysis at the group level. Neuroimage. doi:10.1016/j.neuroimage.2016.05.023

Chen J, Honey CJ, Simony E, Arcaro MJ, Norman KA, Hasson U. 2016. Accessing Real-Life Episodic Information from Minutes versus Hours Earlier Modulates Hippocampal and High-Order Cortical Dynamics. Cereb Cortex. doi:10.1093/cercor/bhv155

Coalson TS, Van Essen DC, Glasser MF. 2018. The impact of traditional neuroimaging methods on the spatial localization of cortical areas. Proc Natl Acad Sci U S A. doi:10.1073/pnas.1801582115

Cole MW, Reynolds JR, Power JD, Repovs G, Anticevic A, Braver TS. 2013. Multi-task connectivity reveals flexible hubs for adaptive task control. Nat Neurosci. doi:10.1038/nn.3470

Corbetta M, Shulman GL. 2002. Control of goal-directed and stimulus-driven attention in the brain. Nat Rev Neurosci 3:201–215. doi:10.1038/nrn755

Dosenbach NUF, Fair DA, Cohen AL, Schlaggar BL, Petersen SE. 2008. A dual-networks architecture of top-down control. Trends Cogn Sci 12:99–105. doi:10.1016/j.tics.2008.01.001

Duncan J. 2010. The multiple-demand (MD) system of the primate brain: mental programs for intelligent behaviour. Trends Cogn Sci 14:172–179. doi:10.1016/j.tics.2010.01.004

Esteban O, Markiewicz CJ, Blair RW, Moodie CA, Isik AI, Erramuzpe A, Kent JD, Goncalves M, DuPre E, Snyder M, Oya H, Ghosh SS, Wright J, Durnez J, Poldrack RA, Gorgolewski KJ. 2019. fMRIPrep: a robust preprocessing pipeline for functional MRI. Nat Methods. doi:10.1038/s41592-018-0235-4

Fedorenko E, Behr MK, Kanwisher N. 2011. Functional specificity for high-level linguistic processing in the human brain. Proc Natl Acad Sci U S A. doi:10.1073/pnas.1112937108

Fedorenko E, Duncan J, Kanwisher N. 2013. Broad domain generality in focal regions of frontal and parietal cortex. Proc Natl Acad Sci U S A 110:16616–16621. doi:10.1073/pnas.1315235110

Ferstl EC, Neumann J, Bogler C, Von Cramon DY. 2008. The extended language network: A meta- analysis of neuroimaging studies on text comprehension. Hum Brain Mapp. doi:10.1002/hbm.20422

Fornito A, Zalesky A, Bullmore ET. 2016. Fundamentals of Brain Network Analysis, Fundamentals of Brain Network Analysis. doi:10.1016/C2012-0-06036-X

Fox MD, Corbetta M, Snyder AZ, Vincent JL, Raichle ME. 2006. Spontaneous neuronal activity distinguishes human dorsal and ventral attention systems. Proc Natl Acad Sci U S A. doi:10.1073/pnas.0604187103

Fruchterman TMJ, Reingold EM. 1991. Graph drawing by force-directed placement. Softw Pract Exp. doi:10.1002/spe.4380211102

Glasser MF, Coalson TS, Robinson EC, Hacker CD, Harwell J, Yacoub E, Ugurbil K, Andersson J, Beckmann CF, Jenkinson M, Smith SM, Van Essen DC. 2016. A multi-modal parcellation of human cerebral cortex. Nature. doi:10.1038/nature18933

Gorgolewski K, Burns CD, Madison C, Clark D, Halchenko YO, Waskom ML, Ghosh SS. 2011. Nipype: A flexible, lightweight and extensible neuroimaging data processing framework in Python. Front Neuroinform. doi:10.3389/fninf.2011.00013

Graesser AC, Hauft-Smith K, Cohen AD, Pyles LD. 1980. Advanced outlines, familiarity, and text genre on retention of prose. J Exp Educ. doi:10.1080/00220973.1980.11011745

Hall P, Wilson SR. 1991. Two Guidelines for Bootstrap Hypothesis Testing. Biometrics. doi:10.2307/2532163

Hassabis D, Maguire EA. 2007. Deconstructing episodic memory with construction. Trends Cogn Sci 11:299–306. doi:10.1016/j.tics.2007.05.001

Hasson U, Chen J, Honey CJ. 2015. Hierarchical process memory: Memory as an integral component of information processing. Trends Cogn Sci. doi:10.1016/j.tics.2015.04.006

Hitchcock D. 2007. Informal Logic and the Concept of ArgumentPhilosophy of Logic. doi:10.1016/B978-044451541-4/50007-5

Hobbes T. 1651. The First Part: Of Man, Chapter III: Of the Consequence or Train of ImaginationLeviathan.

Jacoby N, Fedorenko E. 2018. Discourse-level comprehension engages medial frontal Theory of Mind brain regions even for expository texts. Lang Cogn Neurosci. doi:10.1080/23273798.2018.1525494

James W. 1983. Essays in psychology. Harvard University Press.

Kauppi JP, Jääskeläinen IP, Sams M, Tohka J. 2010. Inter-subject correlation of brain hemodynamic responses during watching a movie: Localization in space and frequency. Front Neuroinform. doi:10.3389/fninf.2010.00005

Kemmerer D. 2014. DiscourseCognitive Neuroscience of Language. Psychology Press.

Knijnenburg TA, Wessels LFA, Reinders MJT, Shmulevich I. 2009. Fewer permutations, more accurate P-valuesBioinformatics. doi:10.1093/bioinformatics/btp211

Lerner Y, Honey CJ, Silbert LJ, Hasson U. 2011. Topographic mapping of a hierarchy of temporal receptive windows using a narrated story. J Neurosci. doi:10.1523/JNEUROSCI.3684-10.2011

Lin N, Yang X, Li J, Wang S, Hua H, Ma Y, Li X. 2018. Neural correlates of three cognitive processes involved in theory of mind and discourse comprehension. Cogn Affect Behav Neurosci. doi:10.3758/s13415-018-0568-6

MacDonald AW, Cohen JD, Andrew Stenger V, Carter CS. 2000. Dissociating the role of the dorsolateral prefrontal and anterior cingulate cortex in cognitive control. Science (80-). doi:10.1126/science.288.5472.1835

Mar RA. 2004. The neuropsychology of narrative: Story comprehension, story production and their interrelation. Neuropsychologia. doi:10.1016/j.neuropsychologia.2003.12.016

Margulies DS, Ghosh SS, Goulas A, Falkiewicz M, Huntenburg JM, Langs G, Bezgin G, Eickhoff SB, Castellanos FX, Petrides M, Jefferies E, Smallwood J. 2016. Situating the default-mode network along a principal gradient of macroscale cortical organization. Proc Natl Acad Sci U S A 113:12574–12579. doi:10.1073/pnas.1608282113

Mineroff Z, Blank IA, Mahowald K, Fedorenko E. 2018. A robust dissociation among the language, multiple demand, and default mode networks: Evidence from inter-region correlations in effect size. Neuropsychologia. doi:10.1016/j.neuropsychologia.2018.09.011

Nastase SA, Gazzola V, Hasson U, Keysers C. 2019. Measuring shared responses across subjects using intersubject correlation. Soc Cogn Affect Neurosci. doi:10.1093/scan/nsz037

Nili H, Wingfield C, Walther A, Su L, Marslen-Wilson W, Kriegeskorte N. 2014. A Toolbox for Representational Similarity Analysis. PLoS Comput Biol 10. doi:10.1371/journal.pcbi.1003553

Oldfield RC. 1971. The assessment and analysis of handedness: The Edinburgh inventory. Neuropsychologia. doi:10.1016/0028-3932(71)90067-4

Park HJ, Friston K. 2013. Structural and functional brain networks: From connections to cognition. Science (80-) 342. doi:10.1126/science.1238411

Petrides M. 1999. Dorsolateral prefrontal cortex: Comparative cytoarchitectonic analysis”in the human and the macaque brain and corticocortical connection patterns. Eur J Neurosci. doi:10.1046/j.1460-9568.1999.00518.x

Power JD, Cohen AL, Nelson SM, Wig GS, Barnes KA, Church JA, Vogel AC, Laumann TO, Miezin FM, Schlaggar BL, Petersen SE. 2011. Functional Network Organization of the Human Brain. Neuron 72:665–678. doi:10.1016/j.neuron.2011.09.006

Power JD, Mitra A, Laumann TO, Snyder AZ, Schlaggar BL, Petersen SE. 2014. Methods to detect, characterize, and remove motion artifact in resting state fMRI. Neuroimage. doi:10.1016/j.neuroimage.2013.08.048

Pruim RHR, Mennes M, van Rooij D, Llera A, Buitelaar JK, Beckmann CF. 2015. ICA-AROMA: A robust ICA-based strategy for removing motion artifacts from fMRI data. Neuroimage. doi:10.1016/j.neuroimage.2015.02.064

Rodero E. 2012. A comparative analysis of speech rate and perception in radio bulletins. Text Talk. doi:10.1515/text-2012-0019

Rumelhart DE. 1975. Notes on a schema for stories,” inRepresentation and Understanding: Studies in Cognitive Science, eds DG Bobrow and A. Collins.

Schacter DL, Addis DR, Buckner RL. 2007. Remembering the past to imagine the future: The prospective brain. Nat Rev Neurosci. doi:10.1038/nrn2213

Schaefer A, Kong R, Gordon EM, Laumann TO, Zuo X-N, Holmes AJ, Eickhoff SB, Yeo BTT. 2018. Local-Global Parcellation of the Human Cerebral Cortex from Intrinsic Functional Connectivity MRI. Cereb Cortex. doi:10.1093/cercor/bhx179

Seeley WW, Menon V, Schatzberg AF, Keller J, Glover GH, Kenna H, Reiss AL, Greicius MD. 2007. Dissociable intrinsic connectivity networks for salience processing and executive control. J Neurosci. doi:10.1523/JNEUROSCI.5587-06.2007

Sepulcre J, Sabuncu MR, Yeo TB, Liu H, Johnson KA. 2012. Stepwise Connectivity of the Modal Cortex Reveals the Multimodal Organization of the Human Brain. J Neurosci 32:10649–10661. doi:10.1523/JNEUROSCI.0759-12.2012

Simony E, Honey CJ, Chen J, Lositsky O, Yeshurun Y, Wiesel A, Hasson U. 2016. Dynamic reconfiguration of the default mode network during narrative comprehension. Nat Commun. doi:10.1038/ncomms12141

Smith M, Milic-Frayling N, Shneiderman B, Mendes Rodrigues E, Leskovec J, Dunne C. 2010. NodeXL: a free and open network overview, discovery and exploration add-in for Excel 2007/2010.

Spreng RN, Mar RA, Kim ASN. 2009. The common neural basis of autobiographical memory, prospection, navigation, theory of mind, and the default mode: A quantitative meta-analysis. J Cogn Neurosci 21:489–510. doi:10.1162/jocn.2008.21029

Thomas Yeo BT, Krienen FM, Sepulcre J, Sabuncu MR, Lashkari D, Hollinshead M, Roffman JL, Smoller JW, Zöllei L, Polimeni JR, Fisch B, Liu H, Buckner RL. 2011. The organization of the human cerebral cortex estimated by intrinsic functional connectivity. J Neurophysiol. doi:10.1152/jn.00338.2011

Vincent JL, Kahn I, Snyder AZ, Raichle ME, Buckner RL. 2008. Evidence for a Frontoparietal Control System Revealed by Intrinsic Functional Connectivity. J Neurophysiol 100:3328–3342. doi:10.1152/jn.90355.2008

Xia M, Wang J, He Y. 2013. BrainNet Viewer: A Network Visualization Tool for Human Brain Connectomics. PLoS One. doi:10.1371/journal.pone.0068910

Xu Y, He Y, Bi Y. 2017. A tri-network model of human semantic processing. Front Psychol. doi:10.3389/fpsyg.2017.01538

Xu Y, Lin Q, Han Z, He Y, Bi Y. 2016. Intrinsic functional network architecture of human semantic processing: Modules and hubs. Neuroimage. doi:10.1016/j.neuroimage.2016.03.004

Zabrucky KM, Moore DW. 1999. Influence of text genre on adults’ monitoring of understanding and recall. Educ Gerontol. doi:10.1080/036012799267440

Zwaan RA, Radvansky GA. 1998. Situation Models in Language Comprehension and Memory. Psychol Bull. doi:10.1037/0033-2909.123.2.162

