## Supplementary Material for "Brain Network Reconfiguration for Narrative and Argumentative Thought"

**Supplementary Figure 1.** Behavior rating on the four selected texts from the participants who participated in the fMRI experiments.

**Supplementary Figure 2.** Distribution of ISC results across brain systems.

**Supplementary Figure 3.** Contrast between the ISC results of narrative and argumentative thought.

**Supplementary Figure 4.** Consistency between the ISC results of two narrative texts.

**Supplementary Figure 5.** Consistency between the ISC results of two argumentative texts.

**Supplementary Figure 6.** Critical functional couplings for narrative and argumentative thought.

**Supplementary Figure 7.** Consistency between the ISFC results of two narrative texts.

**Supplementary Figure 8.** Consistency between the ISFC results of two argumentative texts.

**Supplementary Figure 9.** Data quality check.

**Rating Questionnaire.**

**MRI preprocessing using fMRIPrep 1.50.**

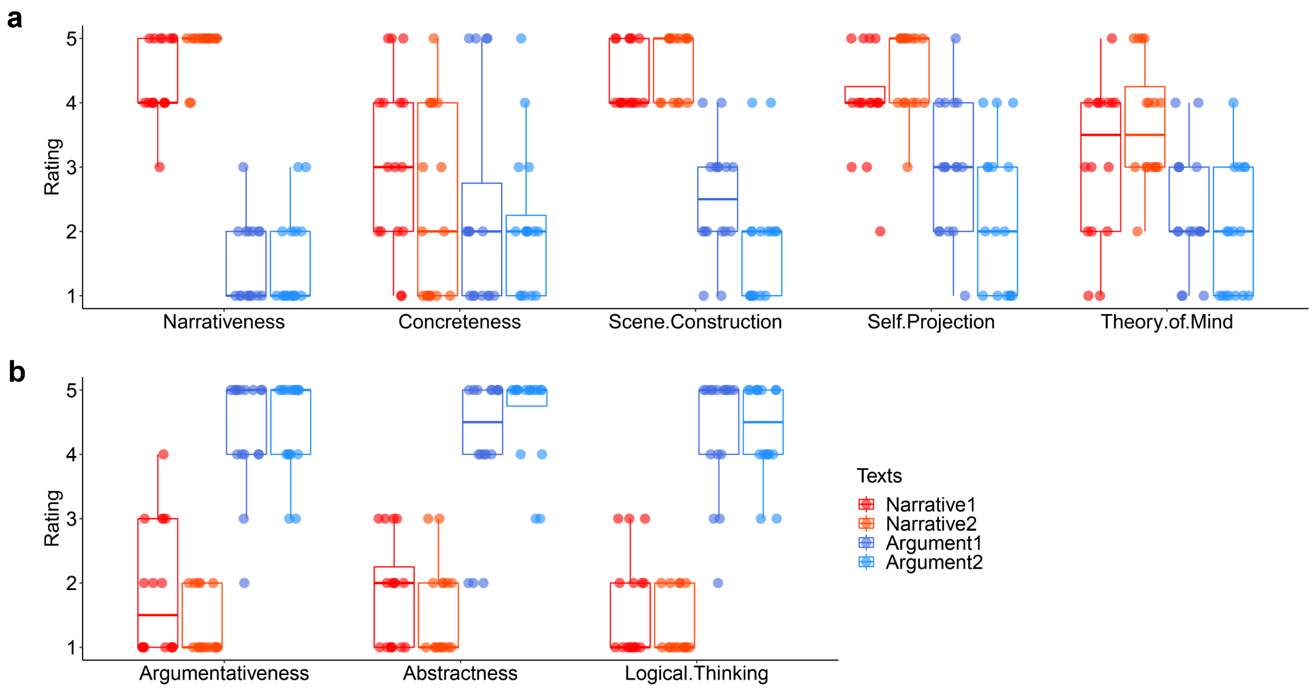


**Supplementary Figure 1. Behavior rating on the four selected texts from the participants who participated in the fMRI experiments.** The boxplots show the rating scores on the four chosen texts from the 16 participants who participated in the fMRI experiment after the scanning. Figure a shows the rating scores on the narrative texts were significantly higher than the argumentative texts on the items of narrativeness (pair $t(15) = 16.37$; $P < 0.001$), scene construction ($pair(15) = 11.28$; $P < 0.001$), self-projection (pair $t(15) = 6.92$; $P < 0.001$), and theory of mind (pair $t(15) = 4.75$; $P < 0.001$). The only exception was the rating on the item of concreteness (pair $t(15) = 1.00$; $P = 0.331$). Figure b shows the rating scores on argumentative texts were significantly higher than the narrative texts on the items of argumentativeness (pair $t(15) = -9.76$; $P < 0.001$), abstractness (pair $t(15) = -9.97$; $P < 0.001$), and logical thinking (pair $t(15) = -10.73$; $P < 0.001$). The results largely validated the rating pattern from an independent group of participants, as shown in Figure 1.

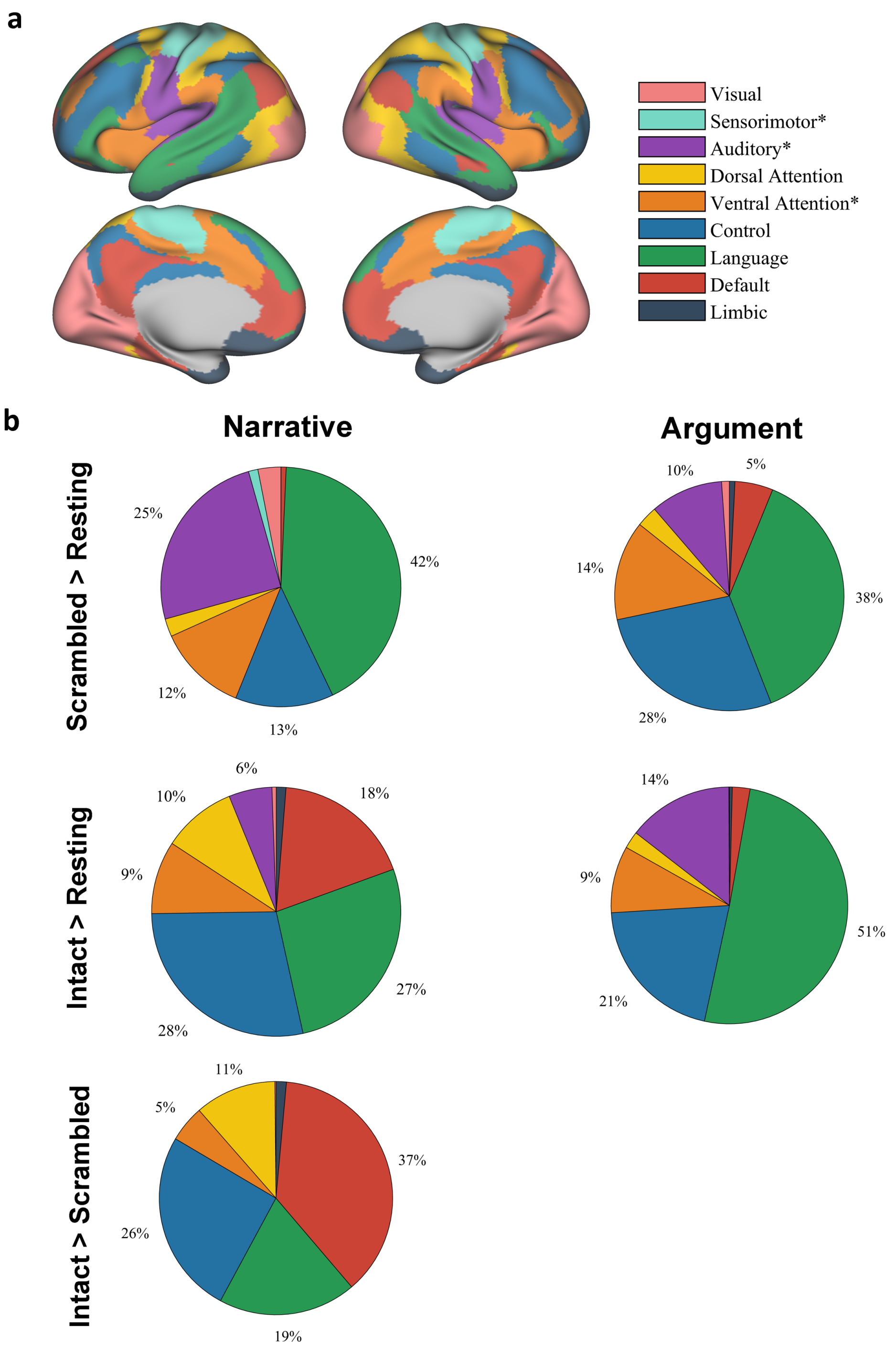


**Supplementary Figure 2. Distribution of ISC results across brain systems.** Figure a shows the spatial distribution of each brain system derived from a pre-labeled atlas (yeo et al., 2011). Figure b shows the pie charts illustrating how many percentages of significant vertexes fall into each brain system in each contrast. The layout corresponds to Figure 2 in the main text. "Sensorimotor*" denotes the brain system including the somatosensory and motor cortices mainly corresponding to the body parts below the neck. "Auditory*" denotes the brain system including not only the auditory cortex but also the somatosensory and motor cortexes mainly corresponding to the body parts above the neck. "Ventral Attention*" denotes the ventral attention network, which may also implicate multiple networks variably referred to in the literature as the salience network or the cingulo-opercular network.

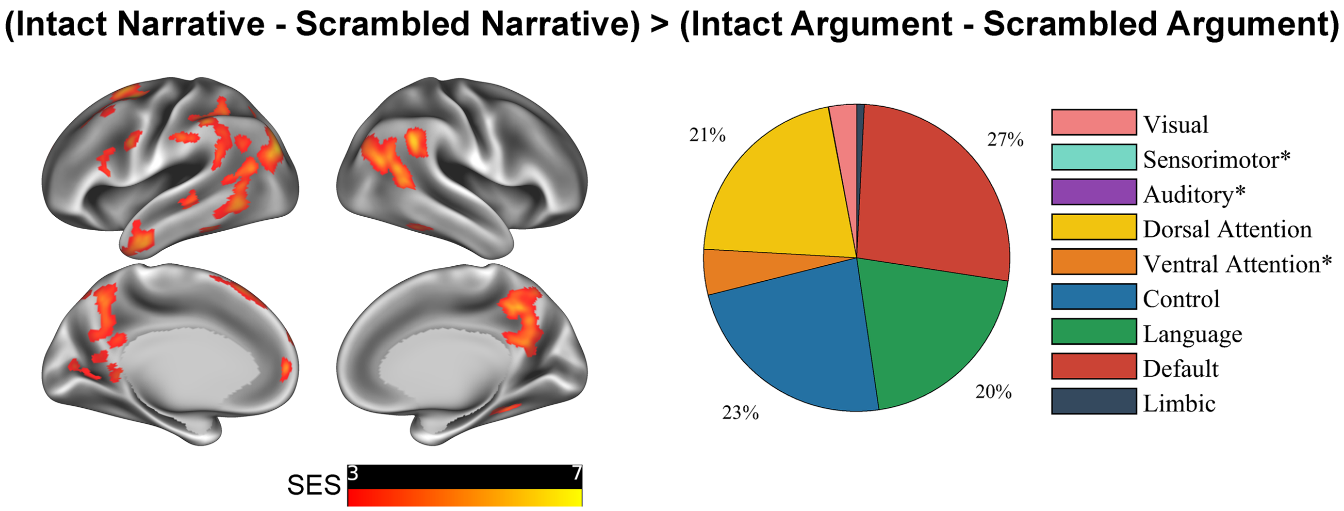


**Supplementary Figure 3. Contrast between the ISC results of narrative and argumentative thought.** The left panel shows the significant areas in the contrast of the ISC results between narrative thought (Intact Narrative – Scrambled Narrative) to argumentative thought (Intact Argument – Scrambled Argument) ($P < 0.05$, FDR corrected, area > 200 $mm^2$). SES = standard effect size. The right panel shows the percentage of the significant areas falling into each brain system. Over 90% of significant brain areas fell into four brain systems. They are dorsal attention, control, language, and default mode. "Sensorimotor*" denotes the brain system including the somatosensory and motor cortices mainly corresponding to the body parts below the neck. "Auditory*" denotes the brain system including not only the auditory cortex but also the somatosensory and motor cortexes mainly corresponding to the body parts above the neck. "Ventral Attention*" denotes the ventral attention network, which may also implicate multiple networks variably referred to in the literature as the salience network or the cingulo-opercular network.

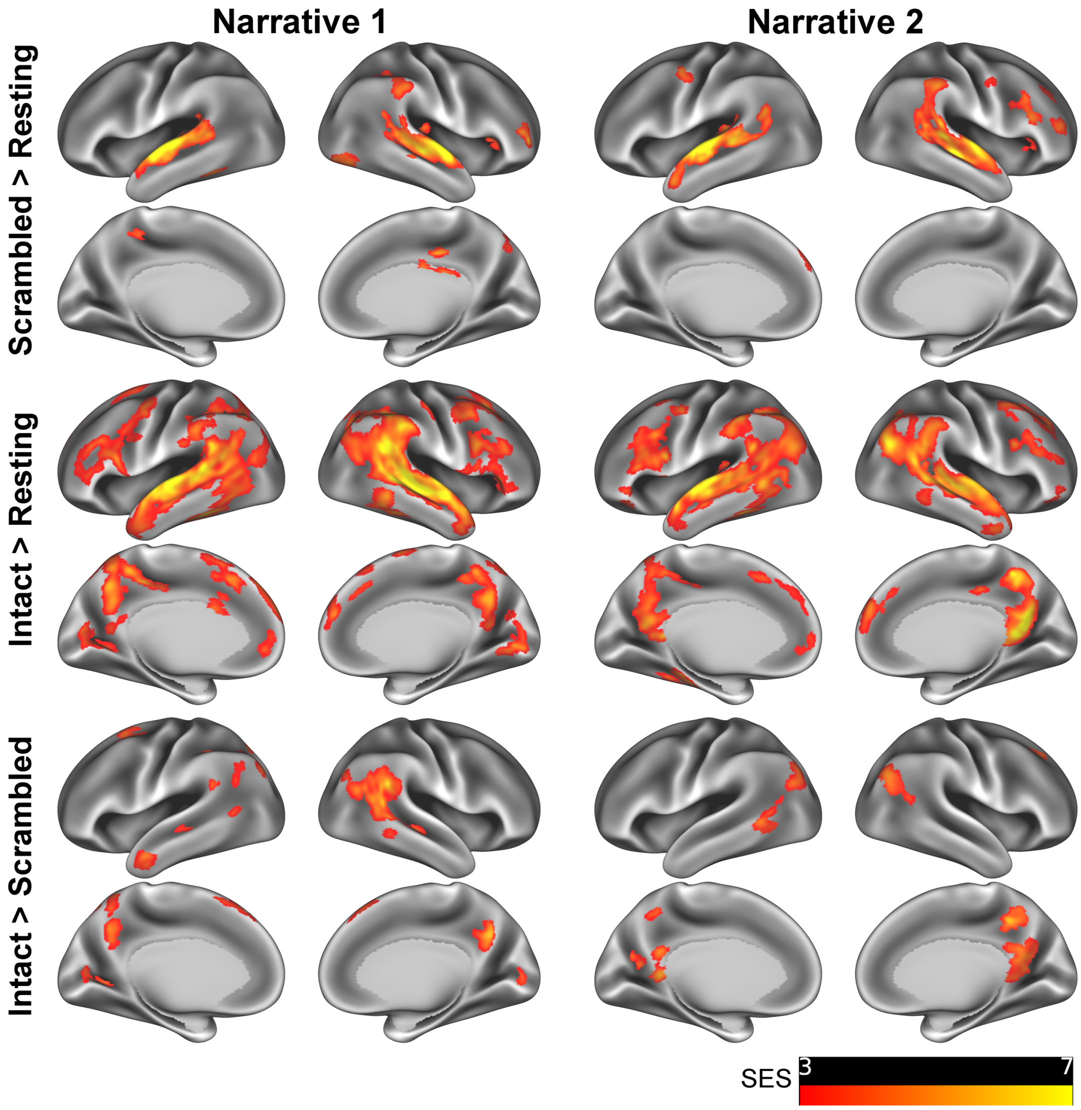


**Supplementary Figure 4. Consistency between the ISC results of two narrative texts.** ISC contrast maps illustrate the significant areas of each contrast using the texts of Narrative 1 (left) and Narrative 2 (right) ($P < 0.05$, FDR corrected, Area > 200 $mm^2$). The results showed an overall consistency between the two narrative texts. The Jaccard similarity coefficients between the two narrative texts in the contrast of the scrambled-sentence condition to the resting state (the first row), the intact-text to the resting state (the second row), and the intact-text condition to the scrambled-sentence condition (the third row) were 0.47, 0.46, and 0.09, respectively. Although the overlap coefficient in contrast between the intact-text condition to the scrambled-sentence condition was relatively low, the intersection area between the two narrative texts mostly fell in the DMN, i.e., the precuneus and the posterior angular gyrus. SES: standard effect size.

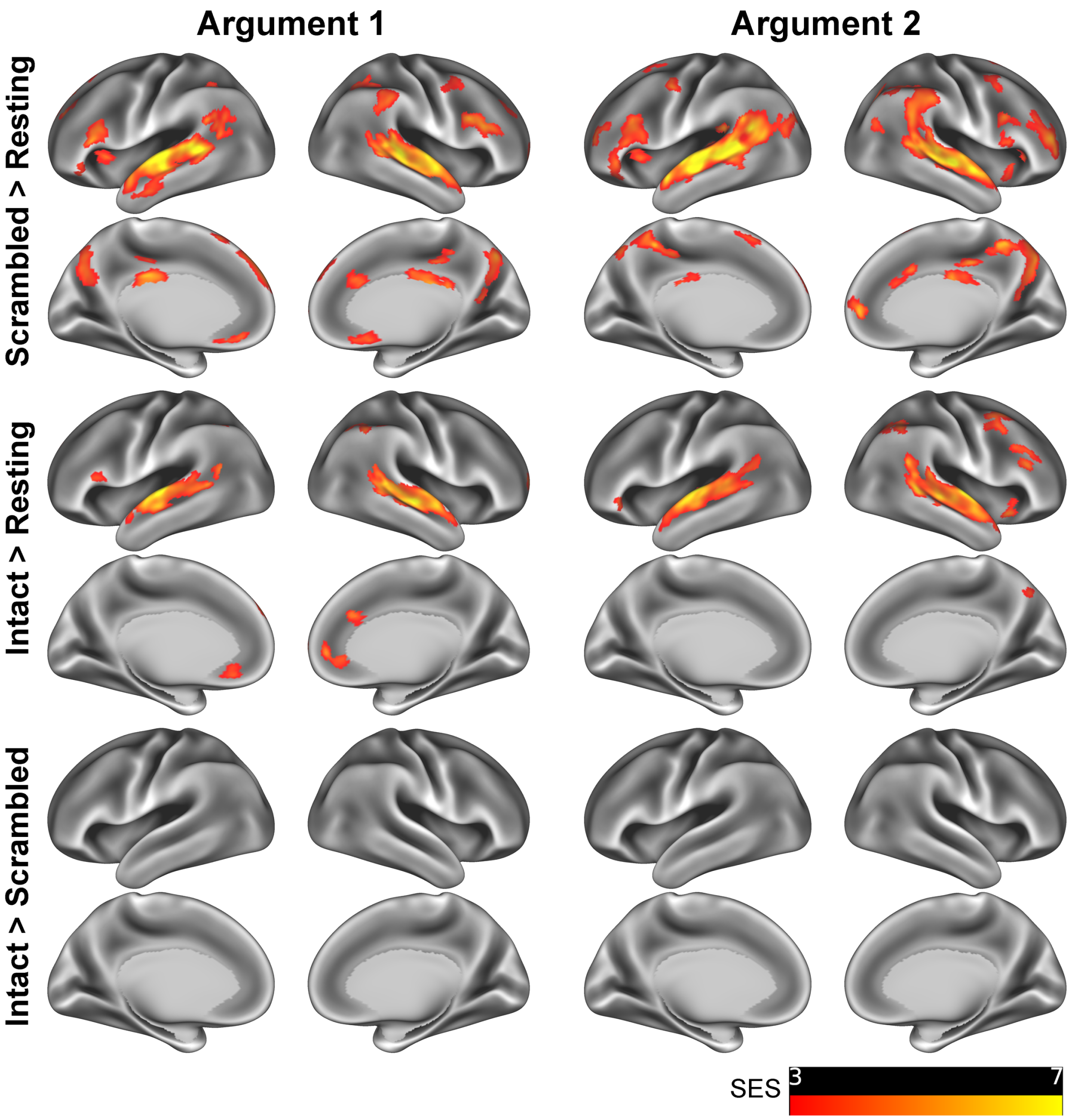


**Supplementary Figure 5. Consistency between the ISC results of two argumentative texts.** ISC contrast maps illustrate the significant areas of each contrast using the texts of Argument 1 (left) and Argument 2 (right) ($P < 0.05$, FDR corrected, Area > 200 $mm^2$). The results showed an overall consistency between the two argumentative texts. The Jaccard similarity coefficients between the two argumentative texts in the contrast of the scrambled-sentence condition to the resting state (the first row) and the intact-text condition to the resting state (the second row) were 0.43 and 0.49, respectively. We did not find any significant areas in the contrast of the intact-text condition to the scrambled-sentence condition for the two argumentative texts (the third row). SES: standard effect size.

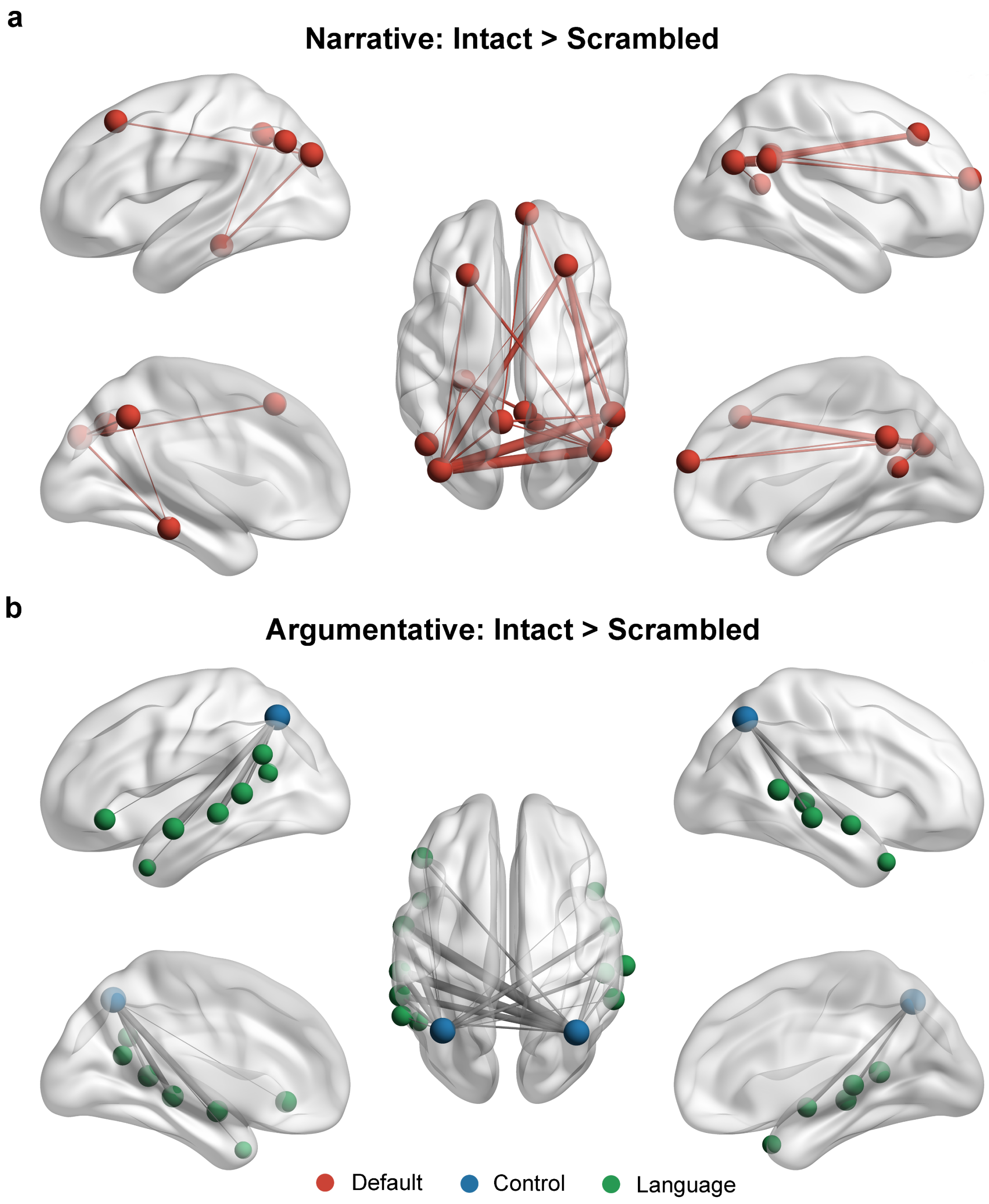


**Supplementary Figure 6. Critical functional couplings for narrative and argumentative thought.** Figure a illustrates the top 20 edges in the DMN with the largest effect size in the contrast between intact-narrative condition and scrambled-narrative condition. All these edges met two criteria: Intact Narrative > Scrambled Narrative ($P < 0.05$, FWE corrected) and Intact Narrative > Resting State ($P < 0.05$, FWE corrected). Figure b illustrates the top 20 edges in the whole brain with the biggest effect size in the contrast between intact-argumentative condition and scrambled-argumentative condition. All these edges met two criteria: Intact Argument > Scrambled Argument ($P < 0.05$, FWE corrected) and Intact Argument > Resting State ($P < 0.05$, FWE corrected). The size of the nodes denotes the node degree in the whole-brain graph. The color of the nodes denotes the brain system that they belong to. The width of the edge denotes the SES of the contrast. Intra-system edges are in the color of that brain system, and inter-system edges are in gray.

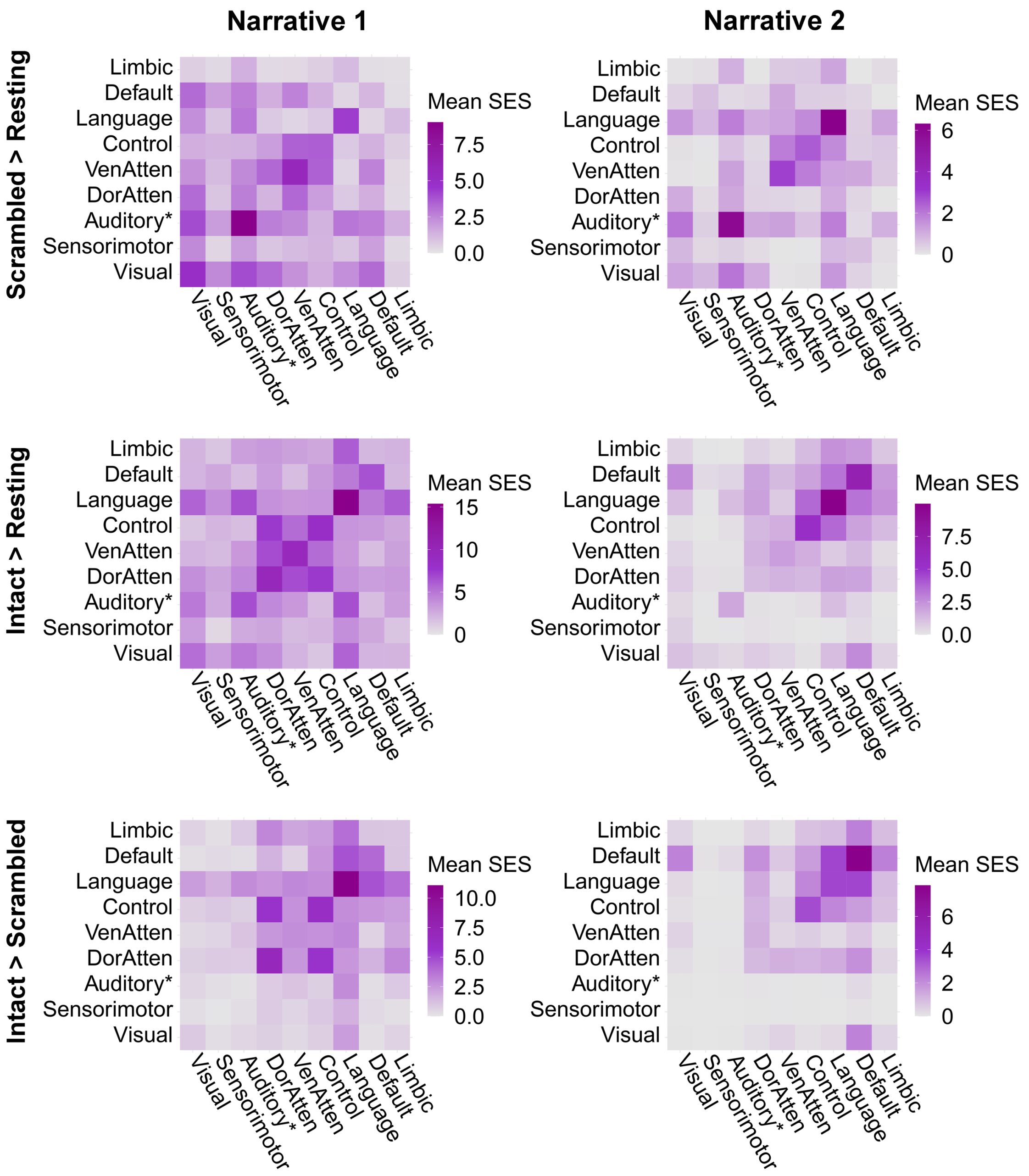


**Supplementary Figure 7. Consistency between the ISFC results of two narrative texts.** The figure compares the network distribution between two narrative texts of all the significant edges ($P < 0.05$, FWE corrected) in the contrasts between the scrambled-narrative condition and the resting state (the first row), the intact-narrative condition and the resting state (the second row), and the intact- and scrambled-narrative conditions (the third row). Each cell indicates the mean SES of each contrast, i.e., the ratio between the sum of the SES and the number of the edges in the fully connected situation. For both of the narrative texts, the scrambled text mainly induced the functional coupling regarding the auditory, language, control, and attention systems. Both intact narratives additionally induced the neural coupling among the brain regions in the DMN. Intact-narrative condition comparing to scrambled-narrative condition enhanced the functional coupling mainly regarding the default mode, language, and control systems. Despite the consistency, different texts of the same type also exhibited a detectable difference in network reconfiguration, which might be due to the variability in content and writing style. "Auditory*" denotes the network including not only the auditory cortex but also the ventral somatosensory and motor brain areas corresponding to the body parts above the neck (see Supplementary Material Fig. 2a). VenAtten = Ventral Attention; DorAtten = Dorsal Attention.

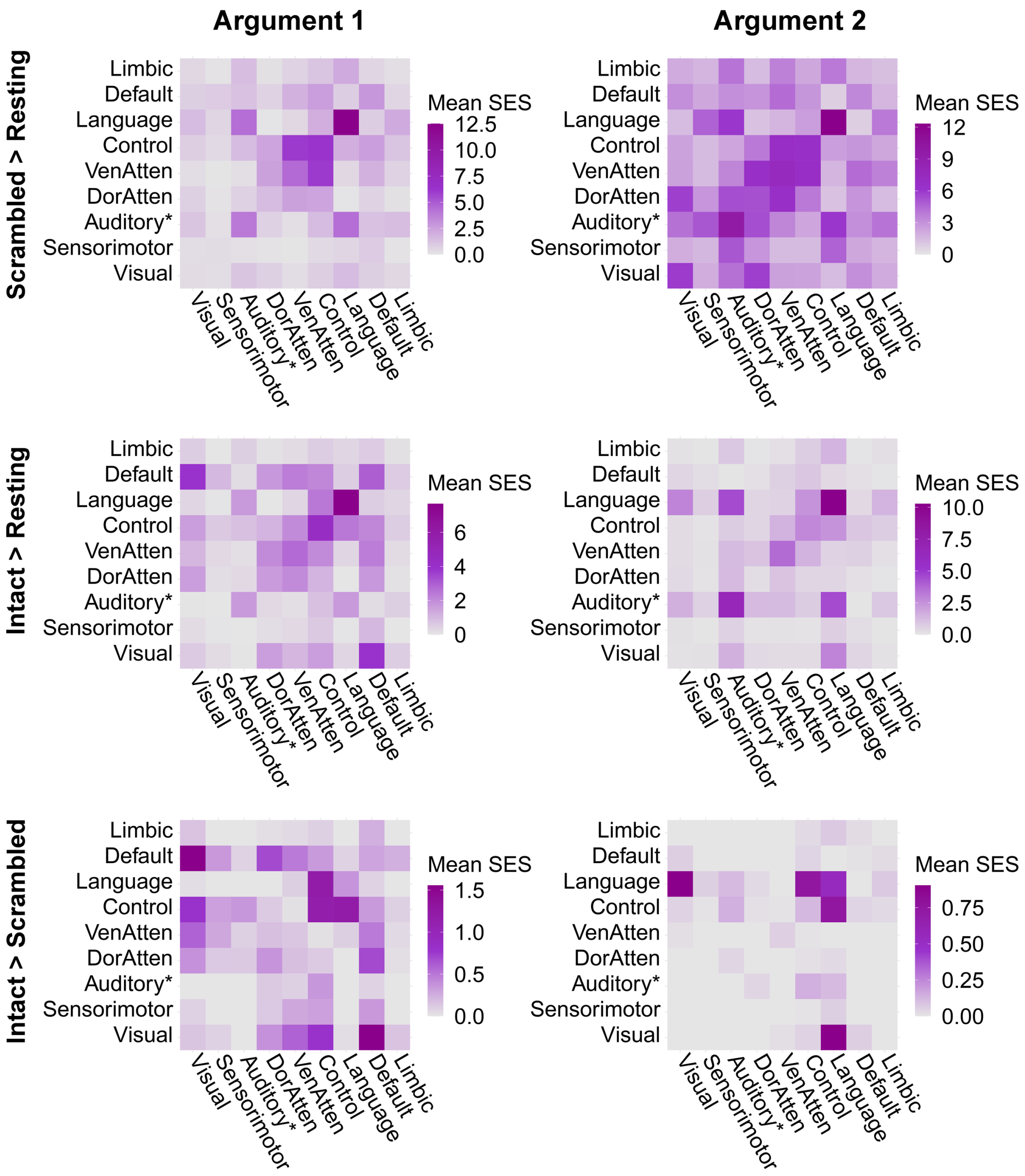


**Supplementary Figure 8. Consistency between the ISFC results of two argumentative texts**. The figure compares the network distribution between two argumentative texts of all the significant edges ($P < 0.05$, FWE corrected) in the contrasts between the scrambled-argumentative condition and the resting state (the first row), the intact-argumentative condition and the resting state (the second row), and the intact- and scrambled-argumentative conditions (the third row). Each cell indicates the mean SES of each contrast, i.e., the ratio between the sum of the SES and the number of the edges in the fully connected situation. For both of the argumentative texts, the scrambled text mainly induced the functional coupling regarding the auditory, language, control, and attention systems. Comparing the intact texts to the scrambled ones revealed the enhanced functional coupling mainly between the control and language systems for both of the argumentative texts. Despite the consistency, different texts of the same type also exhibited a detectable difference in network reconfiguration, which might be due to the variability in content and writing style. "Auditory*" denotes the network including not only the auditory cortex but also the ventral somatosensory and motor brain areas corresponding to the body parts above the neck (see Supplementary Material Fig. 2a). VenAtten = Ventral Attention; DorAtten = Dorsal Attention.

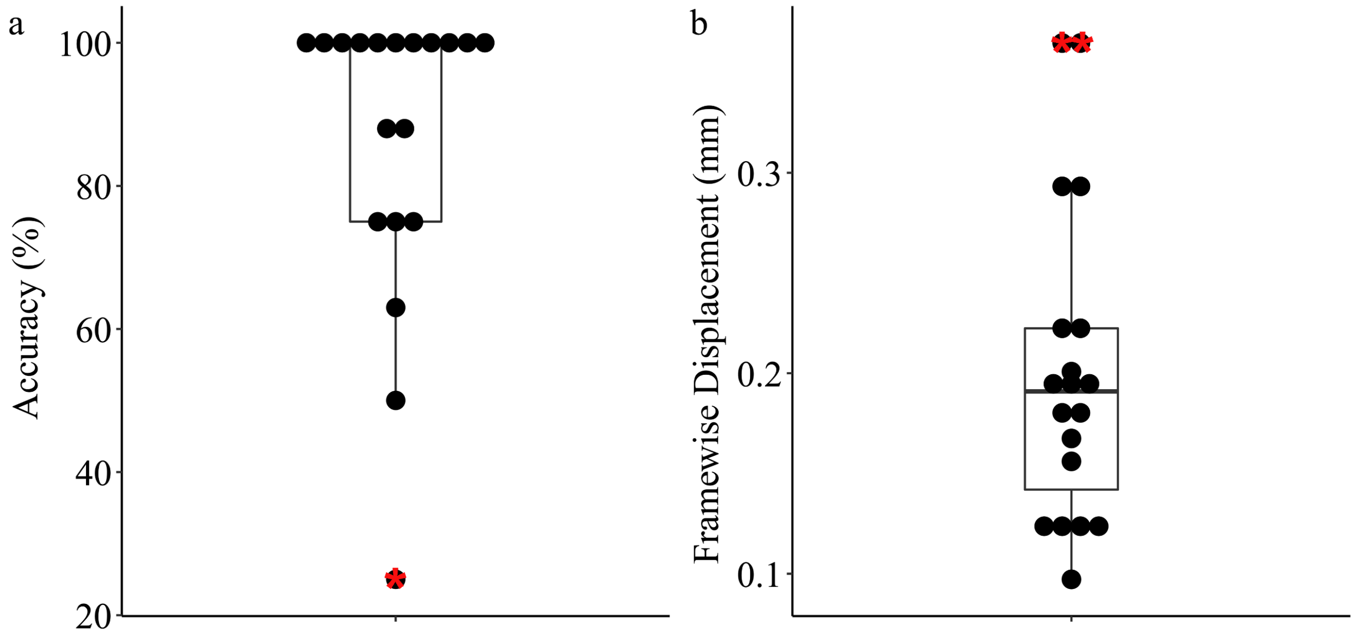


**Supplementary Figure 9. Data quality check.** This figure shows the post-scanning questionnaire results and the head motion measures of the 19 participants. One out of the 20 participants was excluded due to the fuzzy T1 weighted images which cannot be segmented successfully. Figure a illustrates the results of the post-scanning questionnaire. The questionnaire included two questions for each of the four texts regarding its content. The questionnaire aimed to check whether the participants had paid attention to the texts that they heard during the scanning. We excluded one participant whose performance was outside 1.5 times the interquartile range below the lower quartile (denoted with red "*"). Figure b illustrates the head motion of each participant which was measured using the mean frame displacement across nine functional runs. We excluded two participants whose head motion was outside 1.5 times the interquartile range above the upper quartile (denoted with red "*"). The remaining 16 participants entered the analyses.

**Rating Questionnaire**

The questionnaire below shows the English translations of the nine rating questions that were used to select the stimuli. The keywords in the parentheses indicate the features to be rated. They were not presented to the participants. The question order is randomized for each text and for each participant.

Is this text easy to be understood in general? (Difficulty)

1. extremely easy 2. easy 3. not difficult 4 difficult 5. extremely difficult

To what extent do you think this text is a narrative text which tells a story including characters and sequences of events that unfold over time? (Narrative)

1. unlikely 2. hardly likely 3. fairly likely 4. very likely 5. extremely likely

To what extent do you think this text is an argumentative text which claims arguments and shows evidence to persuade the reader to agree with it? (Argumentativeness)

1. unlikely 2. hardly likely 3. fairly likely 4. very likely 5. extremely likely

To what extent do you think this text describes specific experience about a person, a place, or a situation instead of general facts about the universe? (Concreteness)

1. unlikely 2. hardly likely 3. fairly likely 4. very likely 5. extremely likely

To what extent do you think this text exposits general facts, truth, or principles about the universe instead of specific experience? (Abstractness)

1. unlikely 2. hardly likely 3. fairly likely 4. very likely 5. extremely likely

To what extent did the text lead you to imagine the scenes described in the text? (Scene Construction)

1. unlikely 2. hardly likely 3. fairly likely 4. very likely 5. extremely likely

To what extent did the text lead you to mentally put yourself into the situation that describes by this text? (Self-projection)

1. unlikely 2. hardly likely 3. fairly likely 4. very likely 5. extremely likely

To what extent did this text lead you to speculate about the preferences, emotions, or thoughts of other people? (Theory of mind)

1. unlikely 2. hardly likely 3. fairly likely 4. very likely 5. extremely likely

To what extent did the text lead to reason logically? (logical thinking)

1. unlikely 2. hardly likely 3. fairly likely 4. very likely 5. extremely likely

### MRI preprocessing using fMRIPrep 1.50

We performed the fMRI preprocessing using fMRIPrep 1.50. The command line used was: *fmriprep BIDS_DIR OUTPUT_DIR -w WORK_DIR --participant_label PARTICIPANT_LABEL --ignore fieldmaps slicetiming --output-spaces MNI152NLin2009cAsym MNI152NLin6Asym:res-2 fsaverage5 --use-aroma --dummy-scans 10 --cifti-output --write-graph*. The following text is copied from the boilerplate text generated by *fMRIPrep*, which allows for clear and consistent description of the preprocessing steps, aiming to improve experimental reproducibility.

Results included in this manuscript come from preprocessing performed using fMRIPrep 1.5.0 (Esteban, Markiewicz, et al. (2018); Esteban, Blair, et al. (2018); RRID:SCR_016216), which is based on Nipype 1.2.2 (Gorgolewski et al. (2011); Gorgolewski et al. (2018); RRID:SCR_002502).

*Anatomical data preprocessing*

The T1-weighted (T1w) image was corrected for intensity non-uniformity (INU) with N4BiasFieldCorrection (Tustison et al. 2010), distributed with ANTs 2.2.0 (Avants et al. 2008, RRID:SCR_004757), and used as T1w-reference throughout the workflow. The T1w-reference was then skull-stripped with a Nipype implementation of the antsBrainExtraction.sh workflow (from ANTs), using OASIS30ANTs as target template. Brain tissue segmentation of cerebrospinal fluid (CSF), white-matter (WM) and gray-matter (GM) was performed on the brain-extracted T1w using fast (FSL 5.0.9, RRID:SCR_002823, Zhang, Brady, and Smith 2001). Brain surfaces were reconstructed using recon-all (FreeSurfer 6.0.1, RRID:SCR_001847, Dale, Fischl, and Sereno 1999), and the brain mask estimated previously was refined with a custom variation of the method to reconcile ANTs-derived and FreeSurfer-derived segmentations of the cortical gray-matter of Mindboggle (RRID:SCR_002438, Klein et al. 2017). Volume-based spatial normalization to two standard spaces (MNI152NLin2009cAsym, MNI152NLin6Asym) was performed through nonlinear registration with antsRegistration (ANTs 2.2.0), using brain-extracted versions of both T1w reference and the T1w template. The following templates were selected for spatial normalization: ICBM 152 Nonlinear Asymmetrical template version 2009c [Fonov et al. (2009), RRID:SCR_008796; TemplateFlow ID: MNI152NLin2009cAsym], FSL's MNI ICBM 152 non-linear 6th Generation Asymmetric Average Brain Stereotaxic Registration Model [Evans et al. (2012), RRID:SCR_002823; TemplateFlow ID: MNI152NLin6Asym].

*Functional data preprocessing*

For each of the 9 BOLD runs found per subject (across all tasks and sessions), the following preprocessing was performed. First, a reference volume and its skull-stripped version were generated using a custom methodology of fMRIPrep. The BOLD reference was then co-registered to the T1w reference using bbregister (FreeSurfer) which implements boundary-based registration (Greve and Fischl 2009). Co-registration was configured with six degrees of freedom. Head-motion parameters with respect to the BOLD reference (transformation matrices, and six corresponding rotation and translation parameters) are estimated before any spatiotemporal filtering using mcflirt (FSL 5.0.9, Jenkinson et al. 2002). The BOLD time-series, were resampled to surfaces on the following spaces: fsaverage5. Grayordinates files (Glasser et al. 2013), which combine surface-sampled data and volume-sampled data, were also generated. The BOLD time-series (including slice-timing correction when applied) were resampled onto their original, native space by applying a single, composite transform to correct for head-motion and susceptibility distortions. These resampled BOLD time-series will be referred to as preprocessed BOLD in original space, or just preprocessed BOLD. The BOLD time-series were resampled into several standard spaces, correspondingly generating the following spatially-normalized, preprocessed BOLD runs: MNI152NLin2009cAsym, MNI152NLin6Asym. First, a reference volume and its skull-stripped version were generated using a custom methodology of fMRIPrep. Automatic removal of motion artifacts using independent component analysis (ICA-AROMA, Pruim et al. 2015) was performed on the preprocessed BOLD on MNI space time-series after removal of non-steady state volumes and spatial smoothing with an isotropic, Gaussian kernel of 6mm FWHM (full-width half-maximum). Corresponding "non-aggresively" denoised runs were produced after such smoothing. Additionally, the "aggressive" noise-regressors were collected and placed in the corresponding confounds file. Several confounding time-series were calculated based on the preprocessed BOLD: framewise displacement (FD), DVARS and three region-wise global signals. FD and DVARS are calculated for each functional run, both using their implementations in Nipype (following the definitions by Power et al. 2014). The three global signals are extracted within the CSF, the WM, and the whole-brain masks. Additionally, a set of physiological regressors were extracted to allow for component-based noise correction (CompCor, Behzadi et al. 2007). Principal components are estimated after high-pass filtering the preprocessed BOLD time-series (using a discrete cosine filter with 128s cut-off) for the two CompCor variants: temporal (tCompCor) and anatomical (aCompCor). tCompCor components are then calculated from the top 5% variable voxels within a mask covering the subcortical regions. This subcortical mask is obtained by heavily eroding the brain mask, which ensures it does not include cortical GM regions. For aCompCor, components are calculated within the intersection of the aforementioned mask and the union of CSF and WM masks calculated in T1w space, after their projection to the native space of each functional run (using the inverse BOLD-to-T1w transformation). Components are also calculated separately within the WM and CSF masks. For each CompCor decomposition, the k components with the largest singular values are retained, such that the retained components' time series are sufficient to explain 50 percent of variance across the nuisance mask (CSF, WM, combined, or temporal). The remaining components are dropped from consideration. The head-motion estimates calculated in the correction step were also placed within the corresponding confounds file. The confound time series derived from head motion estimates and global signals were expanded with the inclusion of temporal derivatives and quadratic terms for each (Satterthwaite et al. 2013). Frames that exceeded a threshold of 0.5 mm FD or 1.5 standardised DVARS were annotated as motion outliers. All resamplings can be performed with a single interpolation step by composing all the pertinent transformations (i.e. head-motion transform matrices, susceptibility distortion correction when available, and co-registrations to anatomical and output spaces). Gridded (volumetric) resamplings were performed using antsApplyTransforms (ANTs), configured with Lanczos interpolation to minimize the smoothing effects of other kernels (Lanczos 1964). Non-gridded (surface) resamplings were performed using mri_vol2surf (FreeSurfer).

Many internal operations of fMRIPrep use Nilearn 0.5.2 (Abraham et al. 2014, RRID:SCR_001362), mostly within the functional processing workflow. For more details of the pipeline, see the section corresponding to workflows in fMRIPrep's documentation.

*Copyright Waiver*

The above boilerplate text was automatically generated by fMRIPrep with the express intention that users should copy and paste this text into their manuscripts unchanged. It is released under the CC0 license.

*References*

Abraham, Alexandre, Fabian Pedregosa, Michael Eickenberg, Philippe Gervais, Andreas Mueller, Jean Kossaifi, Alexandre Gramfort, Bertrand Thirion, and Gael Varoquaux. 2014. "Machine Learning for Neuroimaging with Scikit-Learn." Frontiers in Neuroinformatics 8. https://doi.org/10.3389/fninf.2014.00014.

Avants, B.B., C.L. Epstein, M. Grossman, and J.C. Gee. 2008. "Symmetric Diffeomorphic Image Registration with Cross-Correlation: Evaluating Automated Labeling of Elderly and Neurodegenerative Brain." Medical Image Analysis 12 (1): 26–41. https://doi.org/10.1016/j.media.2007.06.004.

Behzadi, Yashar, Khaled Restom, Joy Liau, and Thomas T. Liu. 2007. "A Component Based Noise Correction Method (CompCor) for BOLD and Perfusion Based fMRI." NeuroImage 37 (1): 90–101. https://doi.org/10.1016/j.neuroimage.2007.04.042.

Dale, Anders M., Bruce Fischl, and Martin I. Sereno. 1999. "Cortical Surface-Based Analysis: I. Segmentation and Surface Reconstruction." NeuroImage 9 (2): 179–94. https://doi.org/10.1006/nimg.1998.0395.

Esteban, Oscar, Ross Blair, Christopher J. Markiewicz, Shoshana L. Berleant, Craig Moodie, Feilong Ma, Ayse Ilkay Isik, et al. 2018. "FMRIPrep." Software. Zenodo. https://doi.org/10.5281/zenodo.852659.

Esteban, Oscar, Christopher Markiewicz, Ross W Blair, Craig Moodie, Ayse Ilkay Isik, Asier Erramuzpe Aliaga, James Kent, et al. 2018. "fMRIPrep: A Robust Preprocessing Pipeline for Functional MRI." Nature Methods. https://doi.org/10.1038/s41592-018-0235-4.

Evans, AC, AL Janke, DL Collins, and S Baillet. 2012. "Brain Templates and Atlases." NeuroImage 62 (2): 911–22. https://doi.org/10.1016/j.neuroimage.2012.01.024.

Fonov, VS, AC Evans, RC McKinstry, CR Almli, and DL Collins. 2009. "Unbiased Nonlinear Average Age-Appropriate Brain Templates from Birth to Adulthood." NeuroImage 47, Supplement 1: S102. https://doi.org/10.1016/S1053-8119(09)70884-5.

Glasser, Matthew F., Stamatios N. Sotiropoulos, J. Anthony Wilson, Timothy S. Coalson, Bruce Fischl, Jesper L. Andersson, Junqian Xu, et al. 2013. "The Minimal Preprocessing Pipelines for the Human Connectome Project." NeuroImage, Mapping the connectome, 80: 105–24. https://doi.org/10.1016/j.neuroimage.2013.04.127.

Gorgolewski, K., C. D. Burns, C. Madison, D. Clark, Y. O. Halchenko, M. L. Waskom, and S. Ghosh. 2011. "Nipype: A Flexible, Lightweight and Extensible Neuroimaging Data Processing Framework in Python." Frontiers in Neuroinformatics 5: 13. https://doi.org/10.3389/fninf.2011.00013.

Gorgolewski, Krzysztof J., Oscar Esteban, Christopher J. Markiewicz, Erik Ziegler, David Gage Ellis, Michael Philipp Notter, Dorota Jarecka, et al. 2018. "Nipype." Software. Zenodo. https://doi.org/10.5281/zenodo.596855.

Greve, Douglas N, and Bruce Fischl. 2009. "Accurate and Robust Brain Image Alignment Using Boundary-Based Registration." NeuroImage 48 (1): 63–72. https://doi.org/10.1016/j.neuroimage.2009.06.060.

Jenkinson, Mark, Peter Bannister, Michael Brady, and Stephen Smith. 2002. "Improved Optimization for the Robust and Accurate Linear Registration and Motion Correction of Brain Images." NeuroImage 17 (2): 825–41. https://doi.org/10.1006/nimg.2002.1132.

Klein, Arno, Satrajit S. Ghosh, Forrest S. Bao, Joachim Giard, Yrjö Häme, Eliezer Stavsky, Noah Lee, et al. 2017. "Mindboggling Morphometry of Human Brains." PLOS Computational Biology 13 (2): e1005350. https://doi.org/10.1371/journal.pcbi.1005350.

Lanczos, C. 1964. "Evaluation of Noisy Data." Journal of the Society for Industrial and Applied Mathematics Series B Numerical Analysis 1 (1): 76–85. https://doi.org/10.1137/0701007.

Power, Jonathan D., Anish Mitra, Timothy O. Laumann, Abraham Z. Snyder, Bradley L. Schlaggar, and Steven E. Petersen. 2014. "Methods to Detect, Characterize, and Remove Motion Artifact in Resting State fMRI." NeuroImage 84 (Supplement C): 320–41. https://doi.org/10.1016/j.neuroimage.2013.08.048.

Pruim, Raimon H. R., Maarten Mennes, Daan van Rooij, Alberto Llera, Jan K. Buitelaar, and Christian F. Beckmann. 2015. "ICA-AROMA: A Robust ICA-Based Strategy for Removing Motion Artifacts from fMRI Data." NeuroImage 112 (Supplement C): 267–77. https://doi.org/10.1016/j.neuroimage.2015.02.064.

Satterthwaite, Theodore D., Mark A. Elliott, Raphael T. Gerraty, Kosha Ruparel, James Loughead, Monica E. Calkins, Simon B. Eickhoff, et al. 2013. "An improved framework for confound regression and filtering for control of motion artifact in the preprocessing of resting-state functional connectivity data." NeuroImage 64 (1): 240–56. https://doi.org/10.1016/j.neuroimage.2012.08.052.

Tustison, N. J., B. B. Avants, P. A. Cook, Y. Zheng, A. Egan, P. A. Yushkevich, and J. C. Gee. 2010. "N4ITK: Improved N3 Bias Correction." IEEE Transactions on Medical Imaging 29 (6): 1310–20. https://doi.org/10.1109/TMI.2010.2046908.

Zhang, Y., M. Brady, and S. Smith. 2001. "Segmentation of Brain MR Images Through a Hidden Markov Random Field Model and the Expectation-Maximization Algorithm." IEEE Transactions on Medical Imaging 20 (1): 45–57. https://doi.org/10.1109/42.906424.
